# Mesenchymal stromal cells and alpha-1 antitrypsin have a strong synergy in modulating inflammation and its resolution

**DOI:** 10.1101/2022.11.19.517148

**Authors:** Li Han, Xinran Wu, Ou Wang, Xiao Luan, William H. Velander, Michael Aynardi, E. Scott Halstead, Anthony S. Bonavia, Rong Jin, Guohong Li, Yulong Li, Yong Wang, Cheng Dong, Yuguo Lei

## Abstract

Trauma, surgery, and infection can cause severe inflammation. Both dysregulated inflammation intensity and duration can lead to significant tissue injuries, organ dysfunction, mortality, and morbidity. Anti-inflammatory drugs such as steroids and immunosuppressants can dampen inflammation intensity, but they derail inflammation resolution, compromise normal immunity, and have significant adverse effects. The natural inflammation regulator mesenchymal stromal cells (MSCs) have high therapeutic potential because of their unique capabilities to mitigate inflammation intensity, enhance normal immunity, and accelerate inflammation resolution and tissue healing. Furthermore, clinical studies have shown that MSCs are safe and effective. However, they are not potent enough, alone, to completely resolve severe inflammation and injuries. One approach to boost the potency of MSCs is to combine them with synergistic agents. We hypothesized that alpha-1 antitrypsin (A1AT), a plasma protein used clinically and having an excellent safety profile, was a promising candidate for synergism. This investigation examined the efficacy and synergy of MSCs and A1AT to mitigate inflammation and to promote resolution, using in vitro cell cultures and a mouse acute lung injury and inflammation model. We found that the combination of MSCs and A1AT was much more effective than each component alone in i) modulating cytokine releases and inflammatory pathways, ii) inhibiting reactive oxygen species (ROS) and neutrophil extracellular traps (NETs) production by neutrophils, iii) enhancing phagocytosis and, iv) promoting inflammation resolution, tissue healing, and animal survival. Our results support the combined use of MSCs and A1AT for managing severe, acute inflammation.

## INTRODUCTION

Many conditions, including infection, trauma, and surgery, can cause severe inflammation^1–6^. Immune cells are expected to recognize pathogens (or triggers), respond proportionally to the pathogen burden, and effectively eliminate them^7,8^. Subsequently, they initiate a process leading to the resolution of inflammation and restoration of homeostasis^9,10^. Cytokines play critical roles in coordinating immune cell function, ensuring that the initiation, amplification, and resolution of inflammation occurs in an organized manner. Cytokines have a short life span and often remain at the injury site to avoid systemic immune activation. However, under certain conditions, such as an overwhelming pathogen burden, immune cell activation and cytokine production become dysregulated, excessive, persistent, and systemic (i.e., cytokine storm)^1^. Hyperinflammation can rapidly progress to disseminated intravascular coagulation, vascular leakage, acute respiratory distress syndrome (ARDS), multi-organ dysfunction (MODS), and death^11,12^.

Clinical strategies used to treat patients with severe inflammation include supportive care to maintain critical organ functions and elimination of inflammatory stimuli, such as antibiotics. Additionally, steroids and immunosuppressants can be used to suppress immune cells, and targeted biologics (e.g., monoclonal antibodies) can be used to neutralize specific cytokines^1^. However, steroids derail inflammation resolution pathways, compromise antibacterial host defenses, and have significant adverse effects^13–15^. Therefore, there is a clinical need for safe therapies that can mitigate hyper-inflammation while boosting normal immunity and accelerating inflammation resolution.

Our body has multiple types of negative regulators of inflammation, including cells (e.g., Treg)^16^, proteins (e.g., IL-10)^17,18^, and special lipid mediators (e.g., lipoxin A4)^9,13,19–22^. These mechanisms, designed to work together to prevent severe inflammation, often fail in patients with severe medical comorbidities and/or compromised immunity^9,10^. It follows that augmenting these inflammatory regulators may offer a promising therapeutic approach. Among various inflammatory regulators, mesenchymal stromal cells (**MSCs**) are of particular interest since they possess unique and multi-faceted capabilities to mitigate severe inflammation. They can balance the inflammatory environment by downregulating pro-inflammatory cytokines, such as IL-6 and TNFα, while upregulating anti-inflammatory or/and pro-resolving cytokines, such as IL-10 and IL4^23–38^. Using secreted mediators and direct interactions, MSCs can program monocytes and macrophages into the anti-inflammatory and pro-resolving M2 phenotype^24,38^–^41^. They reduce the adherence of leukocytes to endothelium^42^. MSCs can inhibit tissue infiltration as well as ROS and NETs production by neutrophils^24,25,32,35,42–44^. MSCs can also enhance ‘normal’ immunity by boosting the phagocytosis, bacterial killing, and efferocytosis of monocytes and macrophages^39,41,42,45–49^. MSCs also secrete antibacterial peptides such as LL-37, lipocalin-2, and hepcidin^23,28,34,50^. Finally, MSCs can protect organs from inflammation-associated damage while promoting organ healing^25,26,28,29,36,50–54^. MSCs can reduce cell death and improve barrier functions of endothelium and epiththium^24,27,29,30,42,51,54–56^.

In addition to these multiple beneficial functions, MSCs have low immunogenicity. Therefore, allogeneic MSCs can be administered without significant side effects^57^. MSCs can be isolated from various tissues, such as the placenta, umbilical cord, and adipose tissue, and they can be efficiently expanded in vitro. It is therefore hardly surprising that MSCs have been studied in varying disease contexts, including ARDS, sepsis, GvHD, stroke, spinal cord injury, myocardial infarction, organ transplantation, and COVID-19^58–70^. MSCs have also recently been used to treat severe COVID-19 patients^71^, reducing disease mortality significantly^72–76^. However, one shortcoming of MSCs is that monotherapy is not potent enough to fully resolve severe inflammation^77^. Therefore, approaches to boost MSCs’ potency are necessary. One proposed strategy is to combine MSCs with FDA-approved drugs that have excellent safety profiles and can synergize with MSCs.

We propose that protein alpha-1 antitrypsin (**A1AT**) possesses properties well suited to synergize with MSCs and increase their therapeutic efficacy. A1AT is an acute-phase protein whose concentration increases five-fold when the body is injured or infected. A1AT has anti-inflammatory, anti-protease, pro-resolution, cytoprotective, and pro-angiogenic properties^78–88^. It selectively inhibits neutrophil recruitment and cytokine production and neutralizes many pro-inflammatory cytokines^87,89–96^. It suppresses M1 macrophages while promoting M2 macrophages and Treg cells ^78^,^97–104^. It also reduces bacterial and viral burden^105–113^. In addition, it protects cells from various stress^80,114–117^ and promotes angiogenesis^118,119^. A1AT purified from plasma has been used to treat alpha-1 antitrypsin deficiency for decades, with an excellent safety profile^120,121^. Most recently, A1AT has been studied to treat severe COVID-19 patients with positive outcomes^122–126^. However, like MSCs, A1AT alone is insufficient to completely resolve severe inflammation^122–126^. In this investigation, we examined MSCs-A1AT synergism using both in vitro cell cultures and a murine acute lung injury and inflammation model.

## RESULTS

### Isolating MSCs from placenta

The full-term placenta was cut into small pieces, treated with TrypLE for 30 mins, and placed in a cell culture flask (fig. S1A). Cells migrated from the tissues, adhered to the flask surface, and expanded (fig. S1B). When cells reached about 70% confluence, tissues were removed, and cells were allowed to grow until full confluence. These cells were cryopreserved or sub-cultured (fig. S1C). Cells had the classical spindle-like morphology. Above 95% of passage 4 (P4) cells expressed MSC surface markers including CD73, CD90, CD105, CD44, and CD166. The expression of negative markers, including CD45, CD34, CD11b, CD79A, and HLA-DR, was negligible (fig. S1D). In addition, MSCs could be differentiated into FABP4+ adipocytes and osteocalcin+ osteocytes (fig. S1E). In summary, we successfully isolated MSCs from the placenta.

### MSCs modulate cytokine release

To test if our cultured cells could similarly suppress inflammation, we stimulated mouse Raw 264.7 macrophages (MΦs) with LPS and IFNγ to induce intense inflammation. We optimized the concentrations of stimulants, such that 150 ng/mL LPS + 10 ng/mL IFNγ induced maximal cytokine release while not causing rapid and significant cell death. Inflamed cells were treated with MSCs at three different ratios: one MSC for 1, 5, or 10 macrophages (1/1, 1/5, 1/10). 1 μg/mL dexamethasone, a clinically relevant dose used to treat severe inflammation, was used to benchmark MSC’s capability. In addition, one sample was treated with MSCs conditioned medium (CCM) to assess if factors secreted by MSCs were effective. After 24 hrs, the pro-inflammatory (IL6 and TNFα) and anti-inflammatory (IL10) cytokines in the medium were measured with ELISA. The antibodies are specific to mouse proteins to avoid interference from human cytokines secreted by human placenta-derived MSCs.

All treatments reduced the IL6 concentration (fig. S2A). MSCs also decreased TNFα secretion, similar to IL6 (fig. S2B). All treatments except dexamethasone increased IL10 levels. MSCs were better than their conditioned medium (fig. S2C). The IL6/IL10 or TNFα/IL10 ratio can be used to assess inflammation/anti-inflammation balance. Dexamethasone decreased IL6/IL10 from 8 to 3.5, and MSCs decreased IL6/IL10 to 1.5 for 1/10 dosage and to <0.5 for 1/5 and 1/1 dosages. The conditioned medium reduced the ratio to 1.5 (fig.S2D). Dexamethasone decreased TNFα/IL10 from 38 to 18. MSCs decreased TNFα/IL10 to ~5, while the conditioned medium reduced the ratio to ~10 (fig.S2E). In summary, the data showed that i) MSCs could dampen pro-inflammatory cytokine secretion while promoting anti-inflammatory or pro-resolving cytokine secretion; ii) cells were better than their conditioned medium alone and better than dexamethasone; iii) there was no huge difference between the 1/10, 1/5 and 1/1 dose for MSCs in terms of IL6/IL10 or TNFα/IL10 ratios. Thus, we decided to perform subsequent experiments using MSCs at a 1/10 ratio.

### A1AT modulates cytokine release

We evaluated A1AT’s ability to suppress inflammation in Raw 264.7 macrophages. Inflame cells were treated with A1AT (isolated from human plasma) with concentrations ranging from 0.1 to 2.0 mg/mL. A1AT reduced the IL6 and TNFα levels in a dose-dependent manner (fig.S3A, B). A1AT at a concentration ≥0.5mg/mL significantly increased IL10 expression, while dexamethasone did not (fig.S3C). These findings were concordant with previously published data^127^. Dexamethasone decreased IL6/IL10 from 7.5 to 2.2, while A1AT decreased IL6/IL10 to <0.5 when ≥0.5 mg/mL protein was used ^127^. Dexamethasone decreased IL6/IL10 from 7.5 to 2.2, which A1AT decreased IL6/IL10 to <0.5 when ≥0.5 mg/mL protein was used (fig.S3D). Dexamethasone decreased TNFα/IL10 from 30 to 15, while A1AT decreased the ratio to ~2 when the protein was ≥0.5 mg/mL (fig. S3E). In summary, we found that i) A1AT could inhibit pro-inflammatory cytokine secretion while promoting anti-inflammatory/pro-resolving cytokine secretion; ii) there was no significant difference between 0.5, 1.0, and 2.0 mg/mL A1AT in terms of IL6/IL10 or TNFα/IL10 ratios. Therefore, 0.5 mg/mL A1AT was used to perform subsequent experiments.

### MSCs and A1AT have synergy to modulate cytokine release

Next, we studied if MSCs and A1AT exhibited synergistic properties. We treated inflamed Raw 264.7 macrophages with 0.5 mg/mL A1AT alone, 1/10 MSCs alone, or their combination. All treatments reduced IL6 and TNFα levels while increasing IL10 levels, with the MSCs+A1AT combination demonstrating the most significant effect (Fig.1A). Furthermore, we measured 40 inflammation-related cytokines using an antibody array. The treatments affected the expression of 19 cytokines (fig.S4). A1AT reduced the expression of CCL2 (MCP-1), CCL5 (RANTES), CCL17, CXCL1, CXCL9, IFNγ, IL13, IL15, IL1a and IL6 (fig.S4). MSCs reduced the expression of CCL2, CCL17, CXCL9, GM-CSF, IFNγ, IL13, IL15, IL17, IL1a, IL1b, IL6 and TNFα. A1AT and MSCs showed strong synergism in regulating the expression of CCL5, CCL17, CXCL1, CXCL13, CXCL9, G-CSF, GM-CSF, IFNγ, IL10, IL13, IL15, IL1a, IL1b, IL2, IL6, IL7, and TNFα (fig.S4). In summary, the results showed that i) MSCs and A1AT had synergistic effects on regulating many cytokines, and ii) the cytokines affected by A1AT and MSCs were not identical, indicating their mechanisms of action were not identical.

**Fig.1.**
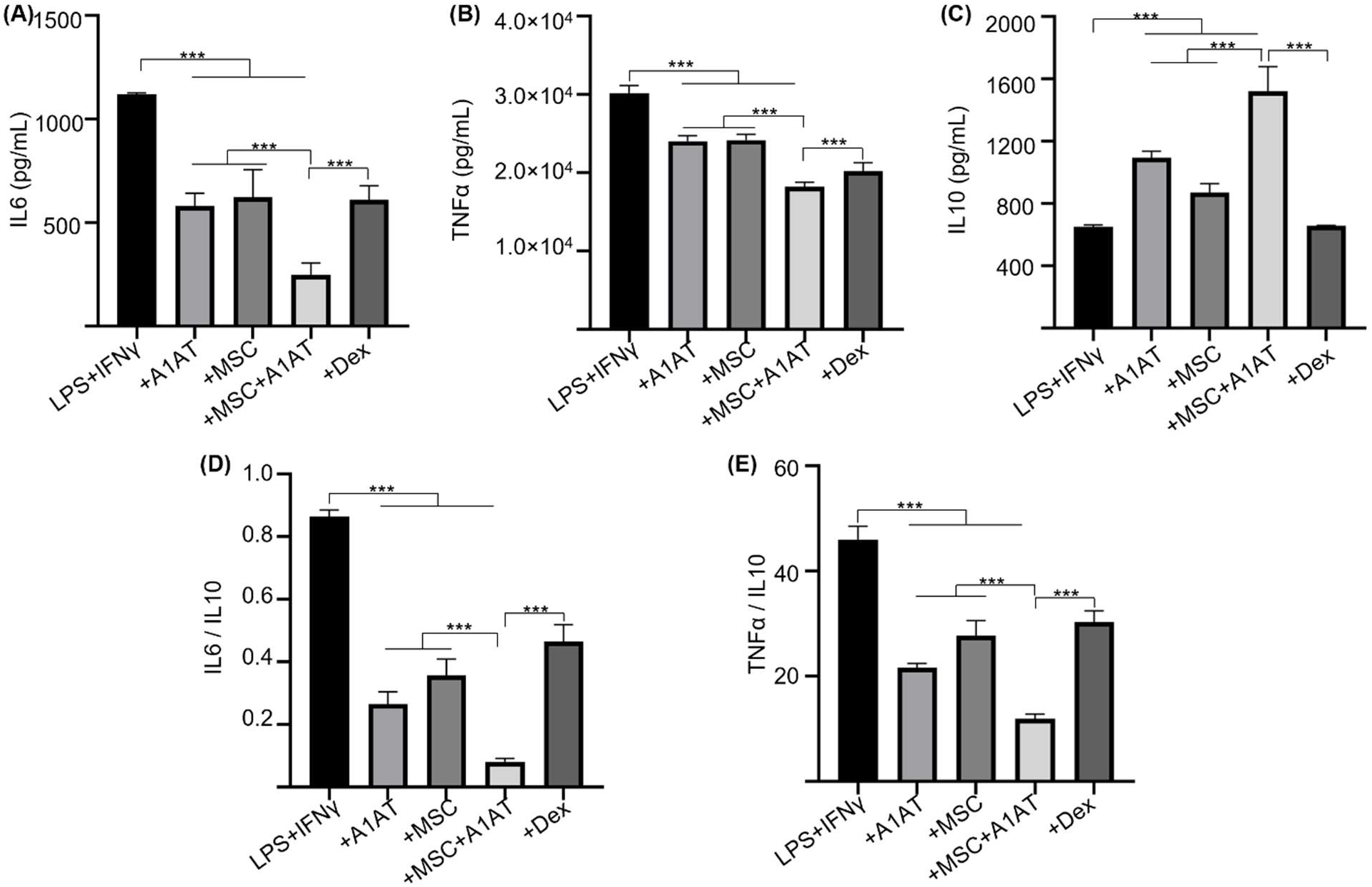
MSCs synergized with A1AT to modulate inflammation in Raw 264.7 macrophages. Cells were stimulated with 100 ng/mL LPS plus 10 ng/mL IFNγ and treated with 0.5 mg/mL A1AT or MSCs (MSC/MΦ=1/10) or their combination. Dexamethasone (Dex, 1 μg/mL) was used as a benchmark. Pro-inflammatory mouse cytokine IL6 (A), TNFα (B), and anti-inflammatory mouse cytokine IL10 (C) were measured via ELISA. The IL6/IL10 (D) and TNFα/IL10 ratio (E) was also shown. *:*p*<0.05, **:*p*<0.01, ***:*p*<0.001.

We then tested whether the findings could be replicated using human macrophages. THP-1 monocytes were first differentiated into macrophages. Inflammation was then induced using LPS and IFNγ. The effects of MSCs, A1AT, and their combination on dampening cytokine release (fig.S5) were similar to Raw 264.7 macrophages (Fig.1). All treatments reduced IL6 and TNFα levels, but only the MSCs+A1AT increased IL10 release. The MSCs and A1AT combination was much more effective than the individual components. The results again showed that MSCs and A1AT could concomitantly downregulate the pro-inflammatory program and upregulate the anti-inflammatory or pro-resolving program.

We also used primary PBMCs to confirm the findings. To avoid donor-to-donor variations, we used PBMCs pooled from multiple donors. We added LPS and IFNγ to activate innate immune cells and anti-CD3 and anti-CD28 antibodies to activate T cells. All treatments reduced IFNγ and TNFα secretion while increasing IL10 production. Again, MSC and A1AT combination was much more effective than the individual components (Fig.2). dexamethasone increased IL10 levels in PBMCs, which is different from the findings using macrophages (Fig.1 and fig.S5). Therefore, we used flow cytometry to assess the cytokine production of monocytes and T cells in PBMCs (fig.S6). Monocytes and T cells were identified with CD14 and CD3 surface markers, respectively. All treatments reduced the % TNFα+ and % IFNγ+ monocytes and their mean fluorescence intensity (fig.S6A). Only MSCs and MSCs+A1AT increased the % IL10+ monocytes and their mean fluorescence intensity. Similar results were found for T cells, except that only MSCs+A1AT increased the % IL10+ monocytes and their mean fluorescence intensity. The results indicated that dexamethasone boosted IL10 production from cell types other than monocytes and T cells in PBMCs.

**Fig.2.**
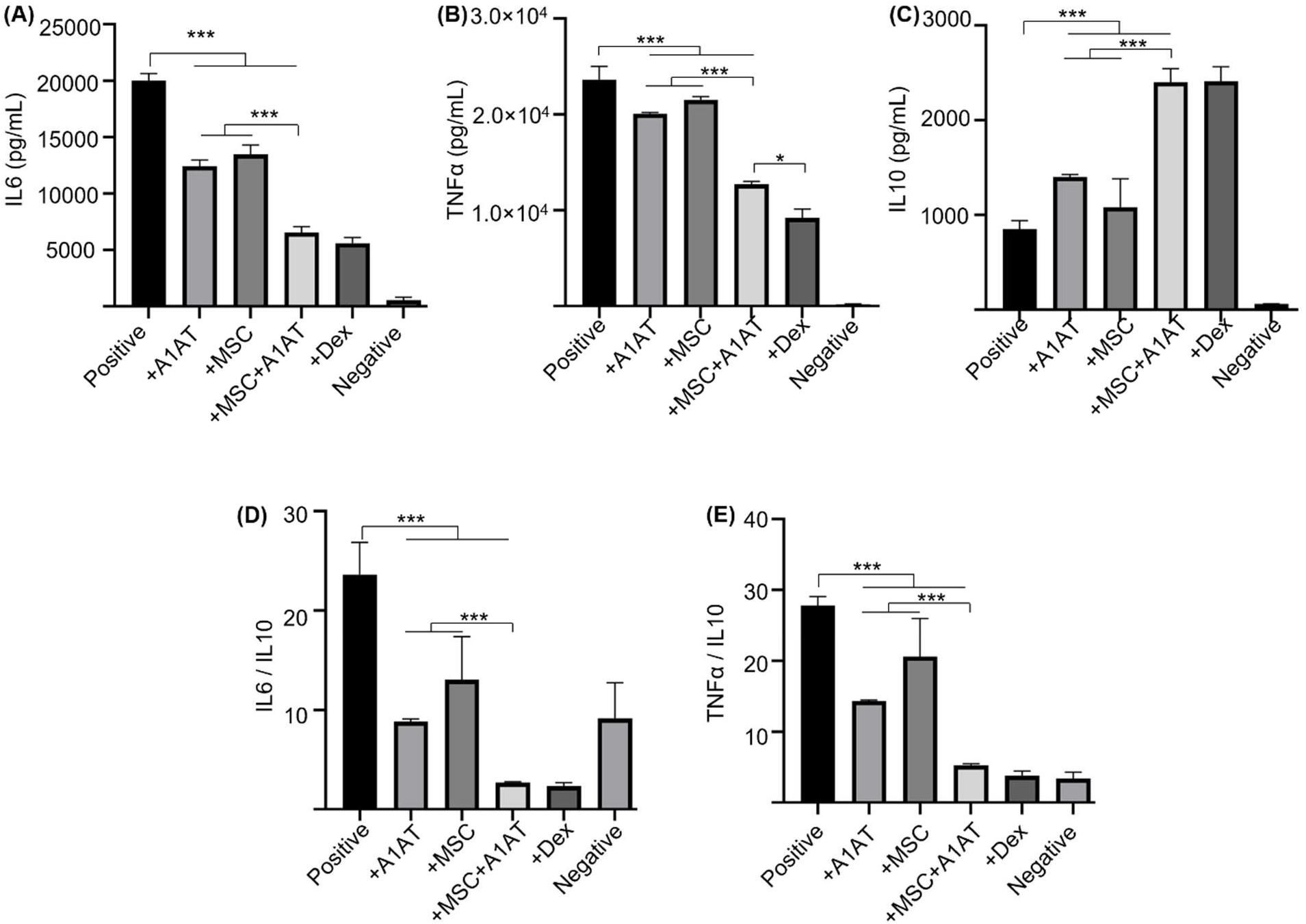
MSCs synergized with A1AT to modulate inflammation in primary human PBMCs. Cells were stimulated with 100 ng/mL LPS + anti-CD3/CD28 antibodies (positive) and treated with 0.5 mg/mL A1AT or MSCs (MSC/PBMC=1/10) or their combination for 24 hrs. Dexamethasone (Dex, 1 μg/mL) was used as a benchmark. PBMCs without activation and treatment were used as a negative control. Pro-inflammatory human cytokine IL6 (A), TNFα (B), and anti-inflammatory human cytokine IL10 (C) were measured via ELISA. The IL6/IL10 (D) and TNFα/IL10 ratio (E) was also shown. *:*p*<0.05, **:*p*<0.01, ***:*p*<0.001.

Furthermore, we measured 40 human inflammation-related cytokines in the PBMCs medium using an antibody array (fig.S7). The treatments affected the expression of 20 cytokines. MSCs reduced the expression of CCL1, CCL5 (RANTES), CXCL13, IFNγ, IL1b, IL2, IL6, IL7 and IL11, while increased IL4 production. A1AT reduced the expression of CCL1, CCL5, CXCL13, CXCL9, G-CSF, CM-CSF, IFNγ, IL12p40, IL1ra, IL1a, IL1b, IL2, IL6, IL7, IL11 and M-CSF, while increased IL10 and IL4 production. A1AT and MSCs showed a strong synergy in regulating the expression of CCL1, CCL5, G-CSF, CM-CSF, IFNγ, IL10, IL12p40, IL1ra, IL1a, IL1b, IL2, IL6, IL7, IL8, IL11, M-CSF and TNFα (fig.S7). The results confirmed the findings using macrophages that i) MSCs synergized with A1AT in regulating many cytokines, and ii) the cytokines affected by A1AT and MSCs were not identical.

### MSCs synergize with A1AT to modulate neutrophil ROS and NETs production

MSCs and A1AT each can inhibit ROS and NETs production^25,128^. We hypothesized that combination therapy would provide synergistic anti-ROS and anti-NET properties when coincubated with neutrophils. Indeed, MSCs+A1AT demonstrated significant synergism in reducing ROS production (Fig.3A, B) and NET production (Fig.3C, D). All treatments also reduced IL6 and TNFα concentrations in the culture medium while increasing the concentration of IL10. In addition, the MSC and A1AT combination worked much better than each treatment alone (fig.S8). In summary, MSCs and A1AT showed a substantial synergy to modulate inflammation and ROS and NETs production in neutrophils.

**Fig.3.**
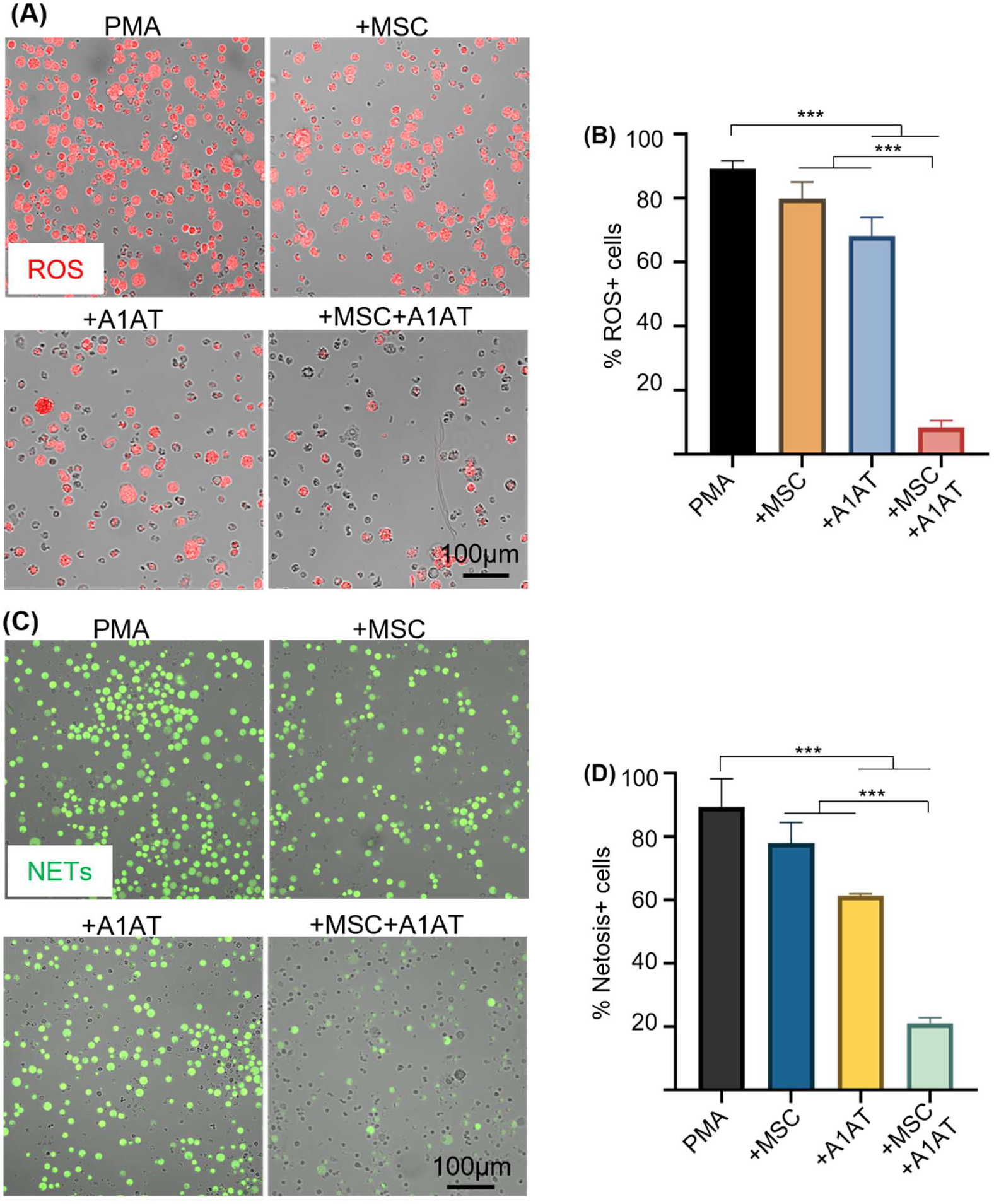
MSCs and A1AT combination treatment reduced neutrophil ROS and NETs production. HL-60 cells derived neutrophils were stimulated with 100 nM PMA and treated with 0.5 mg/mL A1AT or MSCs (MSC/neutrophil=1/10) or their combination for 4 hrs. Reactive oxygen species (ROS) (A, B) and neutrophil extracellular traps (NETs) production (C, D) were analyzed. *:*p*<0.05, **:*p*<0.01, ***:*p*<0.001.

### MSCs synergize with A1AT to modulate macrophage phagocytosis and inflammation pathways

Severe inflammation compromises phagocytosis by innate immune cells, preventing pathogen clearance and inflammation resolution^129–131^. MSCs and A1AT can boost macrophage phagocytosis^38,39,41,42,45–49,100,132^. We thus tested if MSCs and A1AT synergize to enhance phagocytosis in macrophages and neutrophils. We measured the % of cells phagocytosing E. Coli particles, mean fluorescence intensity (MFI) per cell for all cells, and MFI per cell for cells phagocytosing particles. MSCs or A1AT alone did not significantly increase any of these measurements. However, MSCs plus A1AT led to a substantial increase in all these parameters in macrophages (Fig.4 A-D) and neutrophils (Fig.4 E-H).

**Fig.4.**
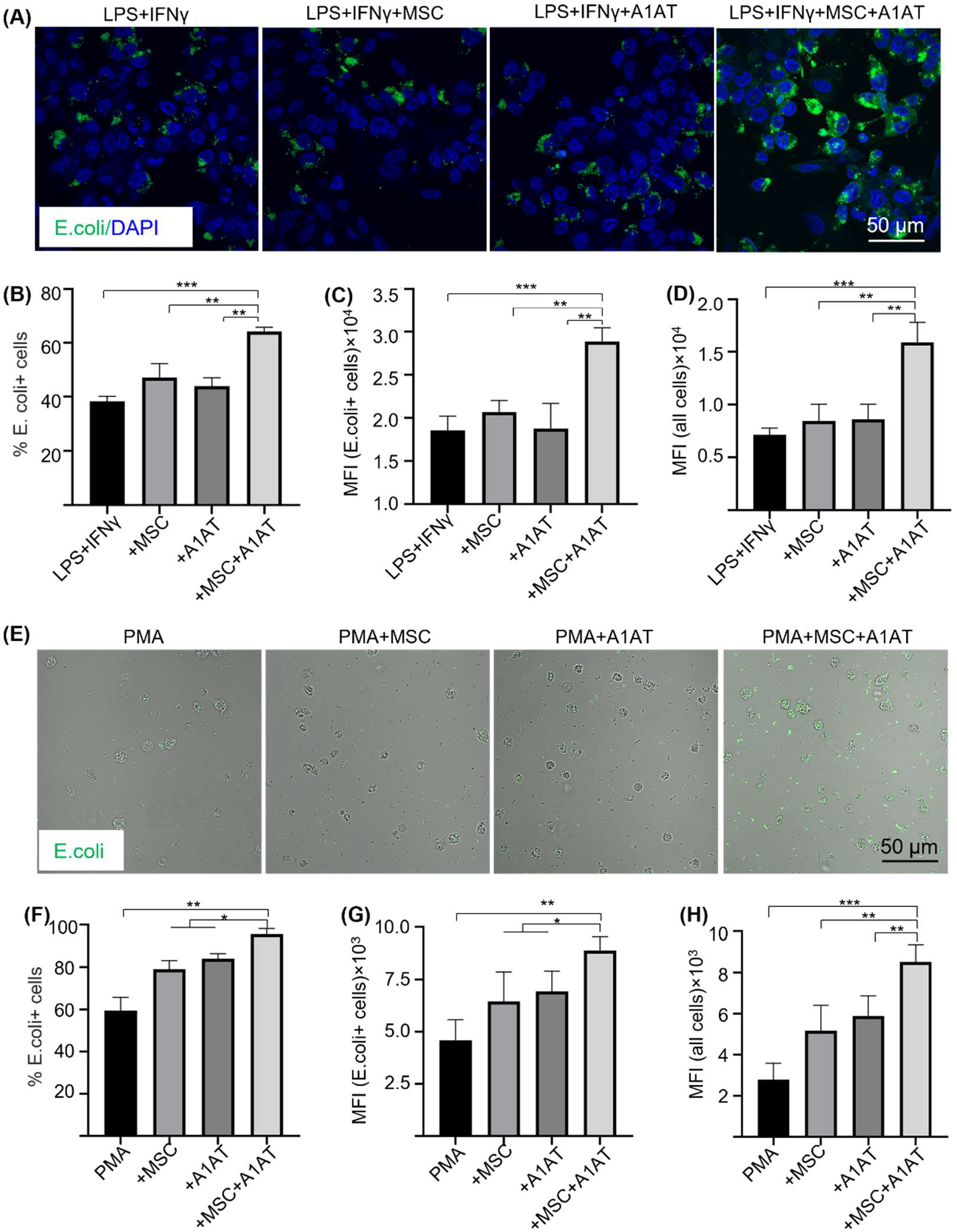
MSC and A1AT combination treatment enhanced phagocytosis in THP-1 derived macrophages (A-D) and HL-60 cells derived neutrophils (E-H). Macrophages were stimulated with 100 ng/mL LPS plus 10 ng/mL IFNγ for 24 hrs. Neutrophils were stimulated with 100 nM PMA for 4 hrs. Cells were treated with 0.5 mg/mL A1AT or MSCs (MSC/MΦ=1/10) or their combination during the stimulation. E. coli particles were added for 3 hrs after treatment. (A, E) E. coli particles emitted green fluorescence after being phagocyted. (B, F) The % E. coli+ cells. (C, G) MFI per cell for all cells. (D, H) MFI per cell for cells with E. coli particles. *:*p*<0.05, **:*p*<0.01, ***:*p*<0.001.

Nuclear factor kappa-light-chain-enhancer of activated B cells (NF-κB) and the interferon regulatory factors (IRF) signaling are critical components of pro-inflammatory pathways. Raw 264.7 and THP-1 cells engineered to express a secreted embryonic alkaline phosphatase (SEAP) reporter for the NF-κB pathway and a secreted luciferase reporter for the IRF pathway were used to evaluate if MSCs and A1AT could regulate these pathways. THP-1 monocytes were differentiated into macrophages before testing. MSCs and A1AT inhibited both pathways in both macrophage types, again demonstrating strong synergistic effects (fig.S9).

### MSCs synergize with A1AT to suppress inflammation and promote inflammation resolution in vivo

We then used the LPS-induced acute lung injury and inflammation mouse model to test if the in vitro results could be replicated in vivo. Treatments were administered 30 mins after the injury (Fig.5A). A lethal dosage (20 mg LPS/kg body weight) was administrated to the first cohort of mice for survival tests. All mice died in 3 days without treatment. MSCs or A1AT alone increased the survival rate, but only their combination wholly protected mice from death (Fig.5B). Furthermore, mice with the combination treatment had significantly less body weight reduction (Fig.5C). A non-lethal dosage (10 mg LPS/kg body weight) was administrated to the second cohort of mice to test inflammation and tissue healing. Tissues were harvested on day 3 for analysis. First, we analyzed lung injury via H&E staining. The lung injury was scored based on five criteria, including i) the number of neutrophils in alveolar space; ii) the number of neutrophils in interstitial space; iii) the amount of hyaline membranes; iv) the amount of proteinaceous debris in airspaces, and v) the alveolar septal thickening. The treatment groups had much less lung injury. The combination therapy group showed the least tissue injury (Fig.5D, E).

**Fig.5.**
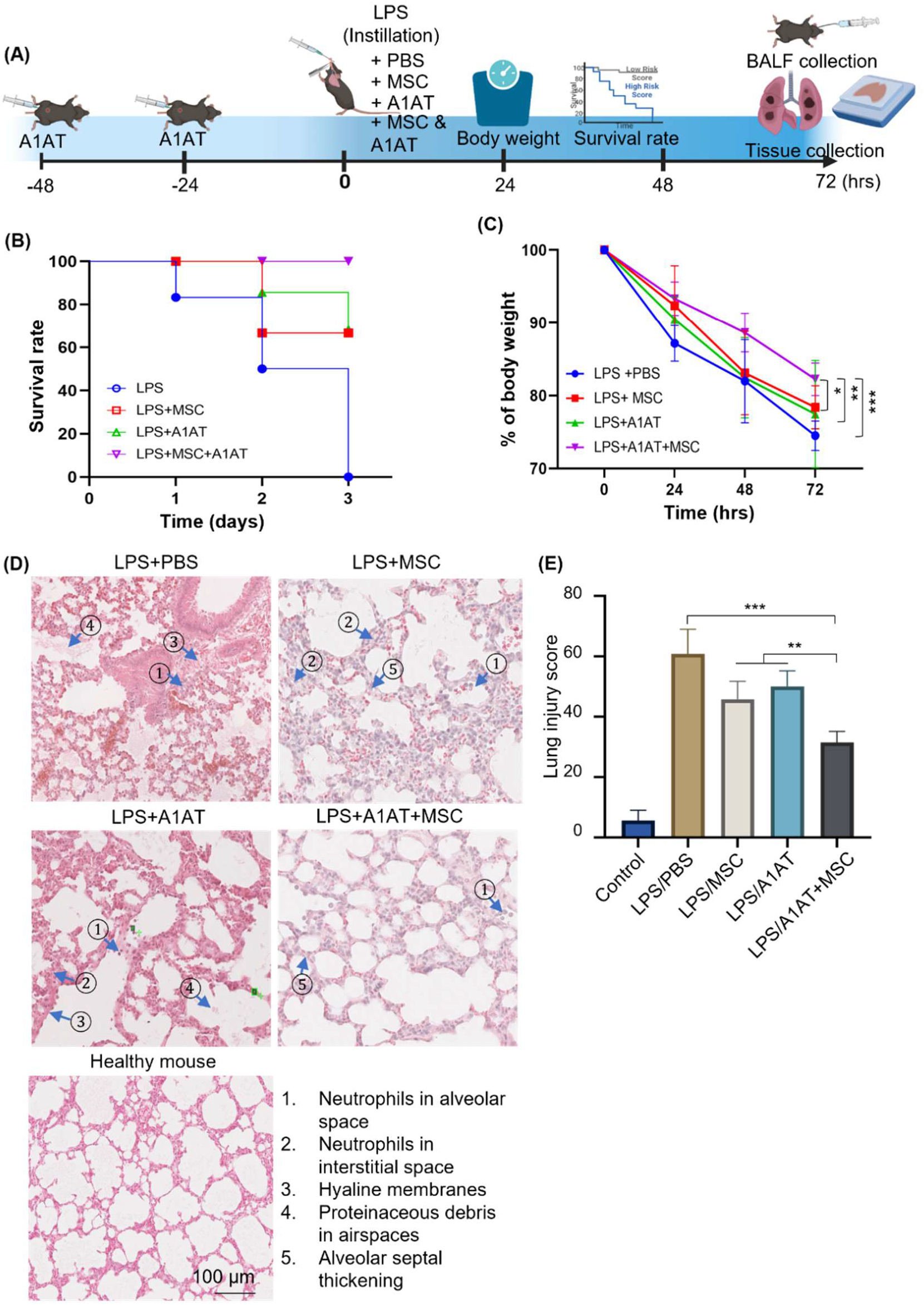
MSCs synergized with A1AT to improve survival rate and reduce lung injury in mice. (A) Illustration of the model. (B) The survival rate and (C) body weight development. n=6. (D) H&E staining and (E) lung injury scores. The lung injury scores were calculated based on the five criteria shown in (D). *:*p*<0.05, **:*p*<0.01, ***:*p*<0.001.

We harvested the bronchoalveolar lavage fluid (BALF) for protein and immune cell analyses. A high total protein concentration indicates the disruption of the endothelium and epithelium. MSCs and A1AT reduced the total protein level, and their combination worked significantly better (Fig.6A). Similar to the in vitro results, MSCs and A1AT reduced IL6 and sTNFαR levels while increasing IL10 levels significantly. Their combination was much more effective than individual components (Fig.6B-F). We measured 40 inflammation-related cytokines with an antibody array. The treatments affected the expression of 21 cytokines. MSCs and A1AT showed a strong synergy on regulating the expression of CCL5, CXCL1, CXCL9, IFNγ, IL10, IL12p70, IL15, IL17, IL1a, IL1b, IL2, IL3, IL4, IL5, IL6, IL7, Leptin and TNFα (Fig.7A). The cytokine array results from BALF (Fig.7A), in vitro mouse macrophages study (fig.S4), and in vitro human PBMCs study (fig.S7) were similar (Fig.7B).

**Fig.6.**
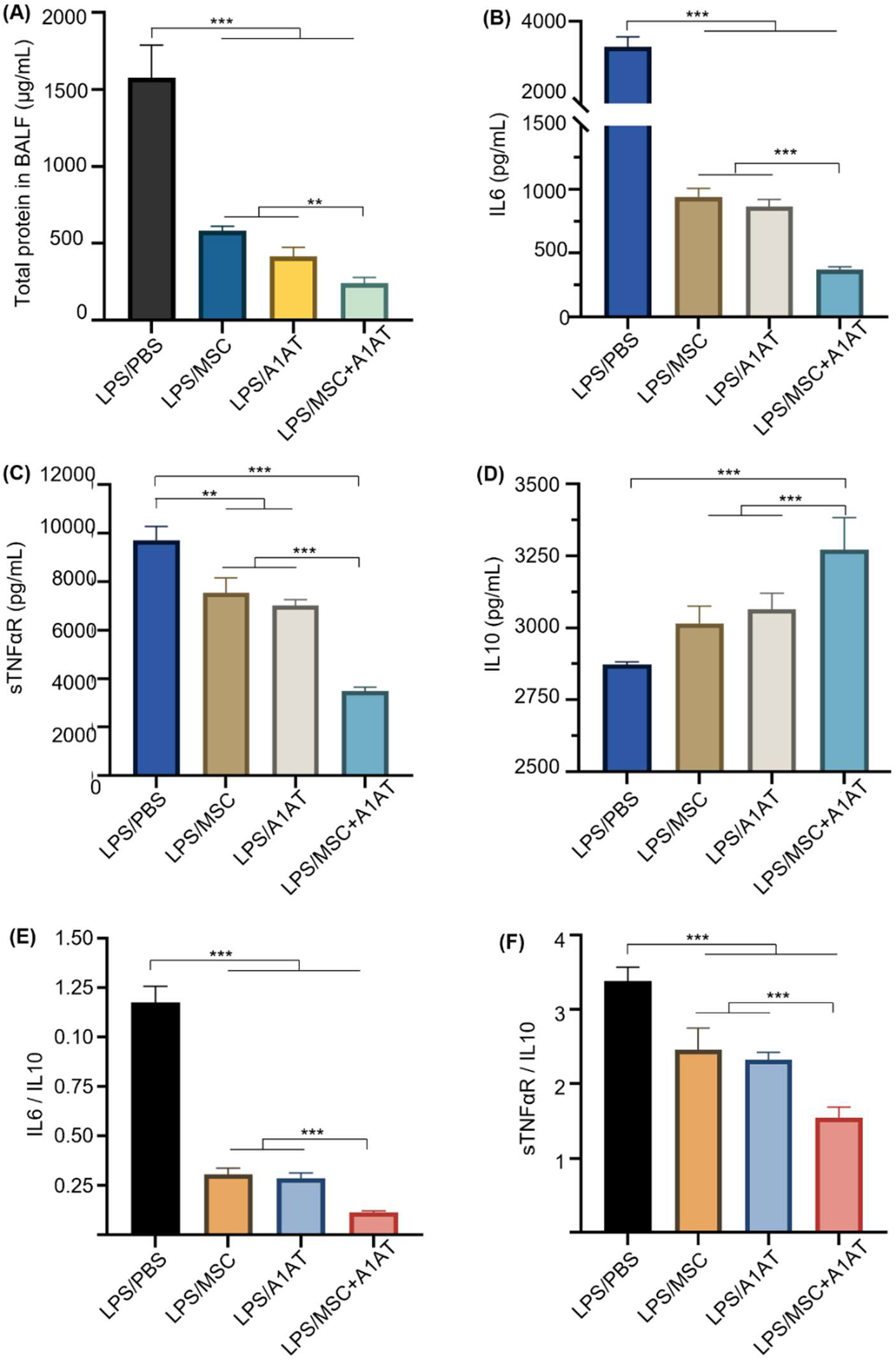
MSCs and A1AT synergized in reducing total protein (A) and pro-inflammatory cytokines while increasing anti-inflammatory cytokine IL10 (B-F) in BALF. *:*p*<0.05, **:*p*<0.01, ***:*p*<0.001.

**Fig.7.**
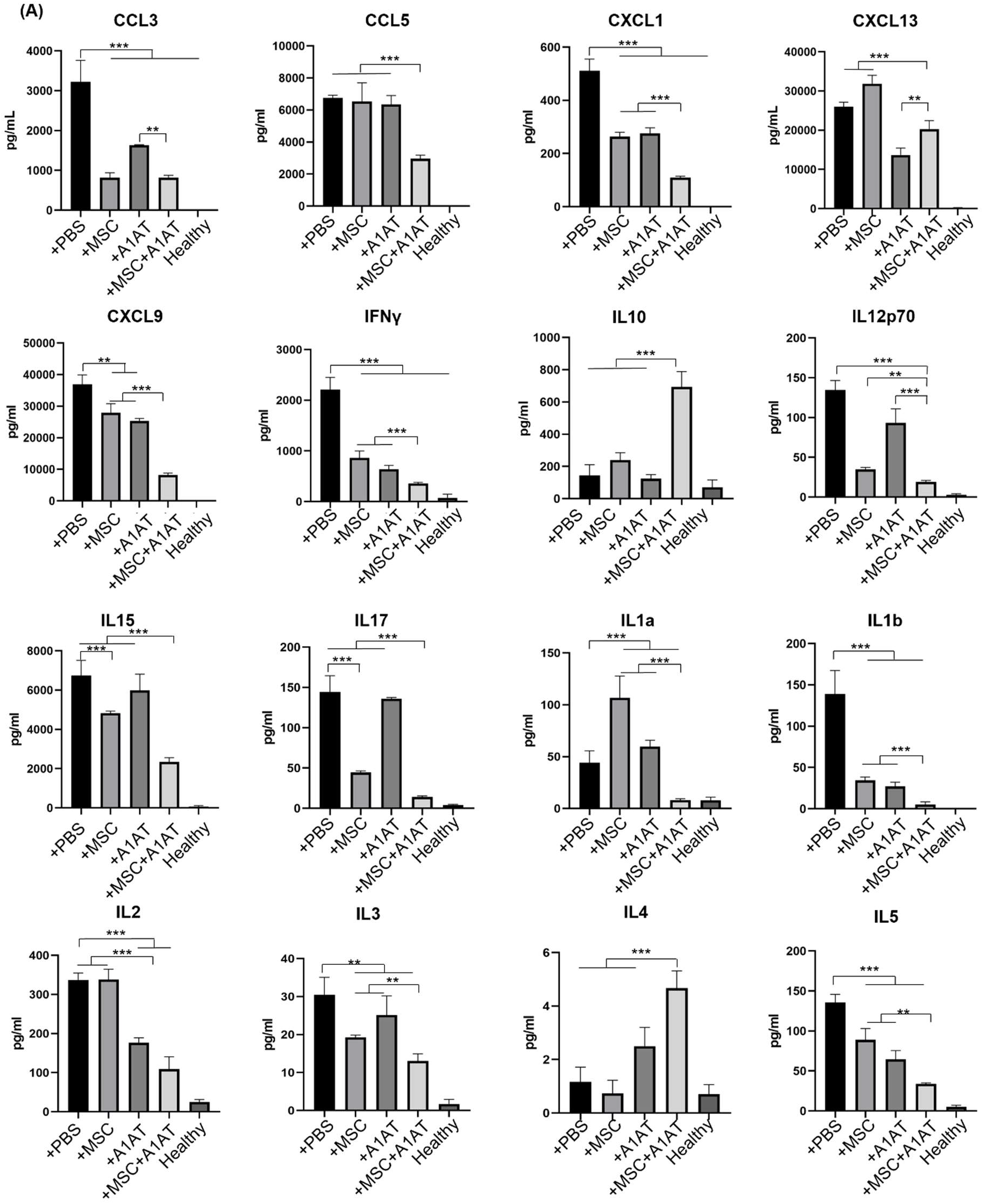
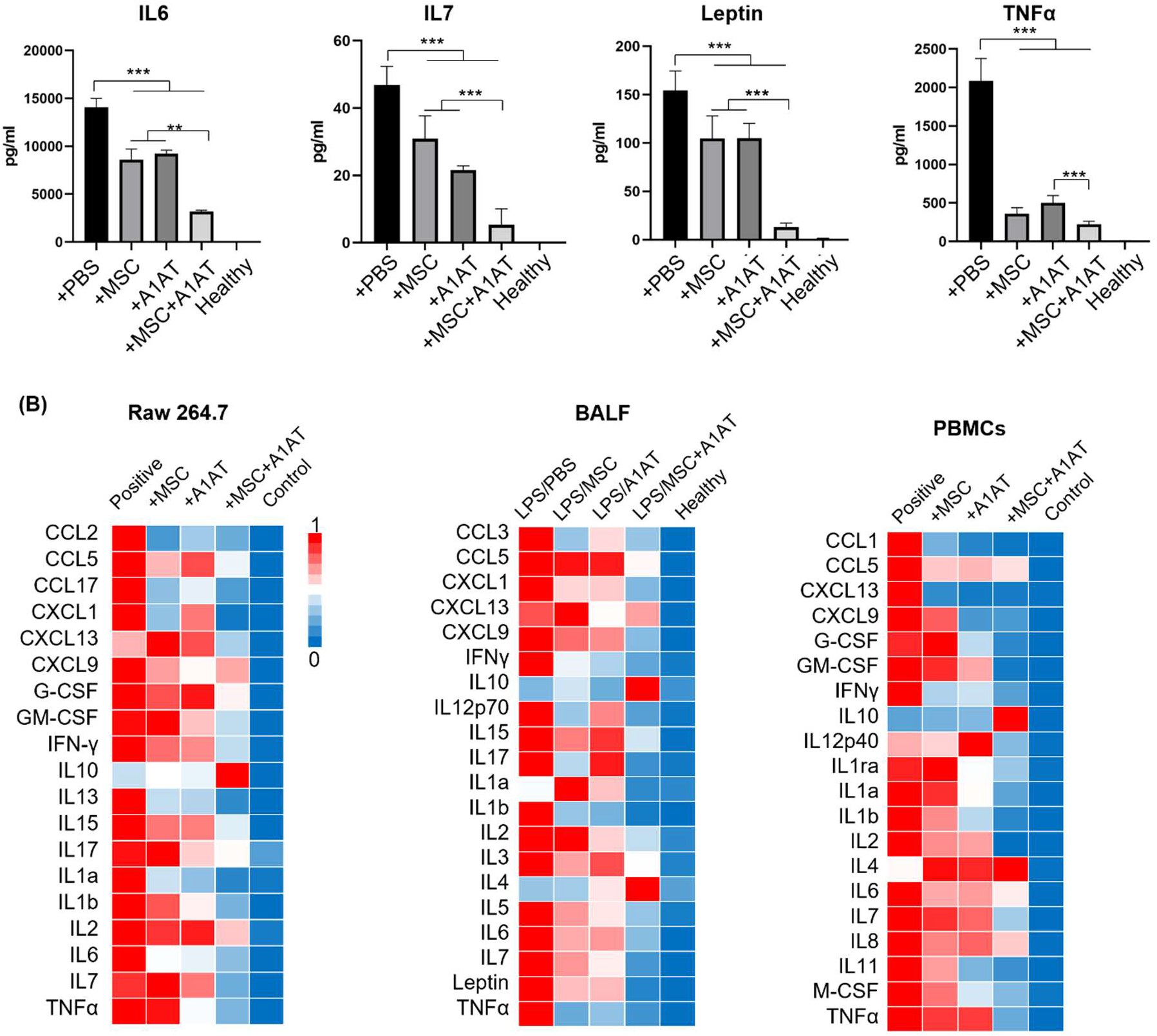
(A) MSC and A1AT combination treatment reduced pro-inflammatory cytokines while increasing anti-inflammatory cytokines in BALF as measured using an inflammation antibody array. Healthy: healthy mouse sample. (B) Heatmaps of cytokine levels in Raw 264.7 medium (from Fig S4), PBMCs medium (from Fig S7), and BALF (from Fig 7a). For each cytokine, the highest expression is set as 1 (red). Other groups are normalized to the highest expression. *:*p*<0.05, **:*p*<0.01, ***:*p*<0.001.

We also analyzed immune cells in BALF. MSCs, A1AT, and especially their combination reduced the number of total cells, macrophages, and neutrophils in BALF. The MSC + A1AT treatment functioned better than the individual components (Fig.8 A-D). The M1/M2 ratio of macrophages was reduced by all treatments (Fig.8E). We used TUNEL staining to identify dead cells in lung tissue. Both MSCs and A1AT reduced the number of dead cells. Dead cells were scarce in the combination treatment group (Fig.8F, G).

**Fig.8.**
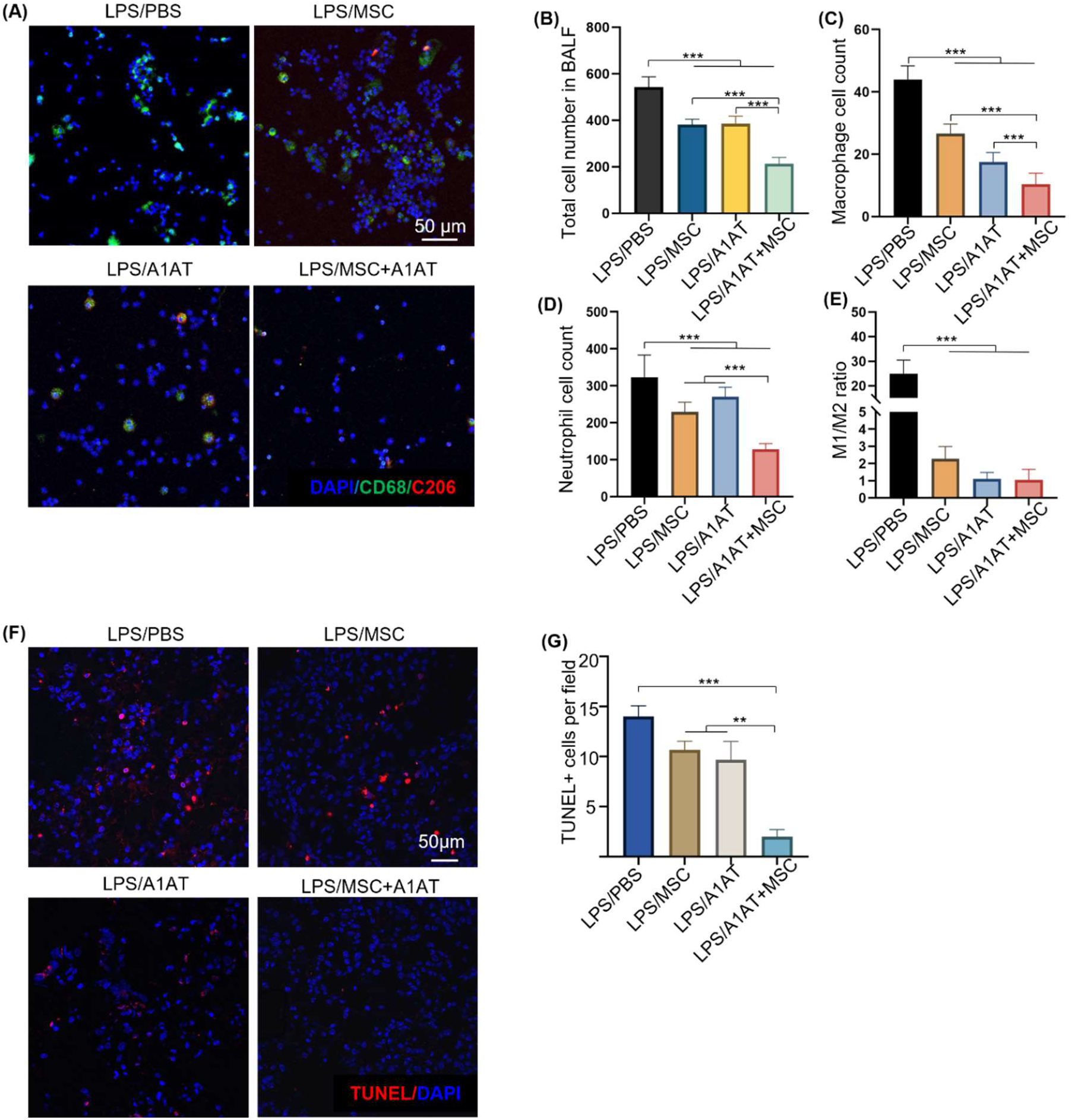
MSCs synergized with A1AT to reduce total cell (A, B), macrophage (C), and neutrophil number (D) in BALF. The M1/M2 macrophage ratio was reduced by all treatments (E). *:*p*<0.05, **:*p*<0.01, ***:*p*<0.001. (F, G) MSCs synergized with A1AT to reduce cell death as identified via TUNEL staining.

## DISCUSSION

Due to their unique ability to mitigate inflammation, boost normal immunity, and promote inflammation resolution and tissue healing, MSCs have been extensively studied in clinical trials for treating severe inflammatory diseases, such as ARDS, sepsis, GvHD, stroke, spinal cord injury, myocardial infarction, multiple sclerosis, organ transplantation, rheumatoid arthritis, Crohn’s, systemic lupus erythematosus, ulcerative colitis and COVID-19^58–70^. A meta-analysis including 55 randomized clinical studies with 2696 patients reported that MSCs induce minor adverse effects while significantly reducing the risk of death^57^. Additionally, no signs of increased tumorgenicity and pro-thrombotic effect were reported^57^. There are about 10 clinical studies on using MSCs to treat ARDS and sepsis^77^. Published results show MSCs are safe and effective in reducing inflammation, epithelial and endothelial damage, and risk of death^60–62,64,67–70,133^. Since the pandemic, >106 registered clinical trials using MSCs to treat severe COVID-19 patients have been initiated^71–74,76,133–140^. Published data show that MSCs can reduce the levels of inflammation biomarkers, pro-inflammatory cytokines, and NETs while increasing the levels of anti-inflammatory cytokines and reducing mortality and morbidity significantly^71,141^. Further, critically ill patients benefitted more from MSC treatment than non-critically ill patients. This finding indicates an additional, unique characteristic of MSCs: they may be able to appropriately respond to the level of inflammation^135^ and are suitable for treating severely ill patients^74^.

A1AT is used to treat alpha-1 antitrypsin deficiency^120,121^. A1AT has also been studied for treating COVID-19^122–126^. Clinical data shows that A1AT concentration is elevated in all COVID-19 patients as a mechanism to counteract inflammation. However, the A1AT response alone is insufficient to resolve the cytokine storm^123^. The IL6/A1AT ratio is significantly higher in severe patients compared to middle patients^123^. A higher IL6/A1AT predicts a prolonged ICU stay and higher mortality^123^. An improvement of IL6/A1AT is associated with better clinical outcomes^123^. A published clinical study finds that A1AT injection can significantly reduce blood IL6 and sTNFR1 levels^125,126^. However, clinical data show that MSCs or A1AT alone are not potent enough to completely resolve hyperinflammation and prevent organ damage^71,125,126,141^. Our data show that MSCs and A1AT demonstrate strong synergy in suppressing pro-inflammatory cytokines, pathways, and NETosis, while boosting anti-inflammatory/pro-resolving factors, normal immunity, and tissue healing. Our study provides strong evidence to support the combined use of MSCs and A1AT for treating severe inflammation in diverse disease states.

A complex network of cells, cytokines, and signaling pathways are involved in hyperinflammation and cytokine storm^1^. Macrophages are major cytokine producers^142–145^. Our data demonstrate that MSCs and A1AT can individually suppress cytokine release from inflamed macrophages and monocytes (Fig.1, 2 and fig.S1-7), confirming previously reported results^24,38–41^. We further demonstrate that combination therapy exceeds the performance of each component (Fig.1, 2, and fig.S1-7). Neutrophils also play a critical role in hyperinflammation^146–155^. Activated neutrophils release NETs and ROS to eradicate bacteria^156^. However, excessive NETs can cause collateral damage to the endothelium, epithelium, and surrounding tissues^157–159^, amplify the cytokine storm^157–159^, and induce disseminated intravascular coagulation^148,160–163^. Our data show that MSCs and A1AT reduce the production of cytokines, ROS, and NETs from neutrophils (Fig.3 and fig.S8), with combination therapy, again exceeding the performance of each individual component. IFNγ release from T cells is crucial to activating macrophages^142–145^. We show that the combination of MSCs and A1AT can significantly suppress TNFα and IFNγ production by T cells (fig.S6). In short, MSCs can synergize with A1AT to effectively modulate the major immune cell types involved in hyperinflammation.

Cytokines IFNγ, IL1, IL6, TNFα, and IL18 play a central role in hyperinflammation^1^. IFNγ is mainly produced by T cells and NK cells and is critical for activating macrophages^142–145^. A recent study finds that IFNγ and TNFα synergistically induce cytokine shock, MODS, and mortality in mice^164^. IL1a/1b bind to IL1 receptors and activate NF-kB to express multiple pro-inflammatory cytokines^165,166^. IL6 acts on both immune and non-immune cells^167–170^. IL6 causes inflammation in endothelial cells, leading to barrier function loss, vascular permeability, hypotension, ARDS, and MODS. TNFα, a potent, multifunctional, pro-inflammatory cytokine, plays a crucial role in a cytokine storm, as shown by the effectiveness of anti-TNF therapies in certain cytokine storm conditions^171–173^. IL10 inhibits the production of TNFα, IL1, IL6, and IL12 and promotes inflammation resolution^174,175^. Our data shows that MSCs synergize with A1AT to simultaneously modulate the major immune cells, cytokines, and pathways involved in severe inflammation (Fig.7 and fig.S4, 7), implying an advantage of this therapy over targeted biologic agents^1^. Neutralizing a particular cytokine with targeted biologics may not always be effective since there is redundancy in pro- and anti-inflammatory pathways^1^.

It should be noted that cytokines modulated by MSCs and A1AT are not identical (Fig 7 and fig.S4, 7), indicating that the cell types and signaling pathways affected by MSCs and A1AT may have differences. This may partly explain their synergism. Our data from mouse macrophages, human macrophages, and PBMCs are congruent in demonstrating the robust efficacy and synergism between MSCs and A1AT (Fig.1–8 and fig.S2-9). Furthermore, the in vivo data agree well with the in vitro results, indicating that the mechanisms of action in vivo can be modeled by the in vitro assays.

The NF-kB pathway plays a pivotal role in inflammation and cytokine storm^176,177^. It can be activated by various ligand-receptor binding such as the binding of LPS to Toll-like receptor 4 (TLR4), the binding of single-stranded viral RNA to TLR7/8 and double-stranded viral RNA to TLR3, and the binding of IL1 and TNFα to their corresponding receptors^176,177^. These lead to the p50/p65 protein translocation to the nucleus to initiate the expression of many pro-inflammatory cytokines, chemokines, adhesion molecules, and growth factors^176,177^. Inhibiting the NF-kB pathway can significantly reduce the cytokine storm, ARDS, MODS, and mortality in animal models with different triggers^176,177^. Glucocorticoids such as dexamethasone and immunosuppressive agents such as Cyclosporin A and tacrolimus are potent NF-kB blockers; however, they have significant adverse effects^178–180^. The IRF pathways also contribute to a cytokine storm. Knocking down the IRF3 and ISGF3 complex in myeloid cells significantly reduces inflammation and mortality in LPS-induced severe inflammation in mice^181,182^. MSCs can inhibit NF-kB signaling^183–186^, which is confirmed by our study. Additionally, we show that the MSCs synergize with A1AT to block both pathways effectively (fig.S9).

An overwhelming pathogen burden often triggers hyperinflammation. Phagocytosis, a major way to clear pathogens, thus represents a valuable therapeutic target to dampen and resolve severe inflammation^130^. Increasing monocytes and macrophage phagocytosis can reduce bacterial burden, cytokine levels, MODS, and mortality^131,187,188^. Clinically, immunoglobulins infused to opsonize and neutralize bacteria, and bacterial products have met modest success^189–192^. G-CSF and GM-CSF have also been studied to increase the neutrophil and macrophage numbers to enhance bacterial clearance with similarly modest success^193–196^. MSCs can boost phagocytosis and bacterial killing of macrophages, thus reducing bacterial burden^39,41,42,45–49^. Our data show that combined MSCs and A1AT can maximally enhance phagocytosis (Fig.4).

Severe inflammation causes ARDS and MODS^1,197–201^. Circulating cytokines upregulate adhesion molecules such as VCAM-1 and ICAM-1 on the endothelium surface while downregulating the tight junction proteins. The adhesion of leukocytes to the endothelium and their trans-endothelium migration is enhanced during severe inflammation. Consequently, large amounts of plasma proteins, cytokines, and immune cells are leaked into parenchymal tissues. They activate the resident immune cells, causing inflammation in distal tissues/organs. The released cytokines and chemokines recruit more immune cells to the tissues. Cytokines, ROS, and proteases cause significant tissue damage. Our data show that MSCs and A1AT reduce BALF’s total protein and immune cells (Fig.8), indicating they can protect the endothelial and epithelial barrier functions. In addition, the total TUNEL+ cells were significantly reduced. Thus, MSCs and A1AT synergize to protect the endothelium, epithelium, and parenchymal tissues. However, since the tissues were harvested 3 days after injury and treatment, the improvement in tissue structure may be because MSCs and A1AT accelerated the inflammation resolution and tissue healing. The higher M2/M1 macrophage ratio and low dead cell number in treatment groups may support this mechanism (Fig.8). Future work should clarify the treatment’s action model and time.

There are a few limitations to the study. First, MSCs and A1AT are only tested in a sterile acute lung injury and inflammation mouse model. Whether the treatment can effectively mitigate severe inflammation caused by infection is unclear, although the features of severe inflammation caused by different triggers are similar. Infection models such as cecal ligation and puncture mice can be used to test the treatment in the future. Testing with large animal models will also be necessary before clinical studies. Second, the molecular mechanisms leading to the MSCs and A1AT synergy are not fully understood. Our data show that MSCs synergize with A1AT to modulate the NF-kB and IFR pathways. We expect there are other pathways contributing to the synergy. Future studies can apply RNA-Seq technology to fully characterize the changes in global gene expressions and signaling pathways caused by the treatments.

In summary, we showed that the MSCs and A1AT combination was much more effective than individual components in i) downregulating pro-inflammatory cytokines while upregulating pro-resolving cytokines, ii) turning off the NF-kB and IRF inflammation pathways, iii) inhibiting neutrophil ROS and NETs production, iv) enhancing macrophage phagocytosis in vitro, and v) reducing the levels of pro-inflammatory cytokines, neutrophils, M1 macrophages, M1/M2 ratio, and tissue injury and mortality significantly in a mouse lung injury model. Our results provide evidence supporting the combined use of MSCs and A1AT as anti-inflammatory therapy. Further investigations are warranted to investigate their combined utility in treating human disease.

## MATERIALS AND METHODS

### Study design

The study was designed to investigate the combinational use of MSCs and A1AT for modulating severe acute inflammation response in vitro and in vivo. All experiments performed in this study had at least three replicates to demonstrate biological reproducibility and to ensure adequate statistical power for comparisons. All animals were randomly allocated to the control and treatment groups. Details for the number of mice, number of cells used, duration, and statistical tests are described below and in the figure legends.

### MSC isolation

Full-term human placentas were purchased from ZenBio Inc. The procedure for isolating and expanding MSCs is similar to a published protocol with minor modifications^7^. Briefly, the placenta was washed and cut into 0.5 cm^3^ pieces that were treated with TrypLE select solution (Gibco) at 37°C for 30 min for partial digestion. 15-20 partially digested pieces were then plated in a 75 cm^2^ tissue flask with 9 mL of EBM-2 complete cell culture medium (EBM-2 +10% FBS+ 1% antibiotic). The flasks were placed in an incubator without disturbance for three days to allow tissues to adhere to the flask surface. After that, the medium was changed every three days until cells reached 70% confluence. These cells were considered passage 0 (P0). They were cryopreserved or subcultured at a seeding density of 5,000 cells/cm^2^ with EBM-2 complete medium.

### MSC surface marker characterization

P4 MSCs were characterized with the Human Mesenchymal Stem Cell Verification Flow Kit (R&D Systems), including antibodies for positive markers CD90, CD73, CD105, and negative markers CD45, CD34, CD11b, CD79A, HLA-DR, as well as the Human Mesenchymal Stem Cells Multi-Color Flow Kit (R&D Systems) including antibodies for positive markers CD44, CD106, CD146, and CD166. Cells were analyzed with the BD FACSCanto™ II System.

### MSC differentiation

P4 MSCs were assessed using the Human Mesenchymal Stem Cell Functional Identification Kit (R&D System) following the product instruction. After 21 days, cells were fixed and stained with FABP-4 antibody to identify adipocytes and osteocalcin antibody to identify osteocytes.

### Immune cell culture

Raw 264.7 cells (RAW-dual cells from InvivoGen) were cultured in DMEM (with 4.5 g/l glucose, 2 mM L-glutamine, 10% heat-inactivated FBS, 100 μg/ml Normocin and 1% Pen-Strep) at a seeding density of 1.5×10^4^ cells/cm^2^. The medium was renewed twice a week. THP-1 cells (THP1-dual cells from InvivoGen) were maintained in RPMI 1640 (with 2 mM L-glutamine, 25 mM HEPES,10% heat-inactivated FBS, 100 μg/ml Normocin, and 1% Pen-Strep). HL-60 cells were cultured in IMEM with 20% FBS.

### Macrophage inflammation assay

Raw 264.7 cells were stimulated with 100 ng/mL LPS (O111:B4, Sigma) plus 10 ng/mL murine IFNγ (Peprotech). Human M0 macrophages were differentiated from THP1 monocytes by incubating cells with 100 ng/mL PMA (Sigma) for 24 hrs. Macrophages were then stimulated with 100 ng/mL LPS plus 10 ng/mL human IFNγ. For treatment, A1AT was added to the medium, and P4 MSCs were co-cultured with macrophages. Condition medium was harvested after 18 hrs, and cytokines were measured by ELISA. The quantitative levels of 40 mouse (for Raw 264.7 and BALF) or human (for PBMCs) cytokines were evaluated with the Mouse or Human Inflammation Arrays (RayBiotech) following the product instructions. Array scanning and data extraction were done by RayBiotech using InnoScan 700/710 Microarray Scanner (Innopsys).

### Neutrophil ROS production

HL-60 cells were differentiated into neutrophil-like cells with 0.1 μM ATRA and 1.25% DMSO in RPMI1640 (with 10% FBS and 2 mM L-Glutamine) for 5 days. Cells were preloaded with 5 μM CellROX deep red reagent (Invitrogen) for 15 min at 37°C. After washing, cells were resuspended in fresh medium and seeded into 96-well plates (100 μL of 200,000 cells/mL/well). Next, cells were activated with 100 nM PMA and treated with 0.5 mg/mL A1AT or 1/10 MSCs or their combination. The fluorescent and phase contrast images were taken with an FV3000 confocal laser scanning microscope (Olympus).

### Neutrophil NETs production

The Incucyte Cytotox Red Dye was used to measure NETs production. HL-60 cells were differentiated into neutrophil-like cells with 0.1 μM ATRA and 1.25% DMSO in RPMI1640 (with 10% FBS and 2 mM L-Glutamine) for 5 days. Cells were preloaded with Cytotox Red Dye and seeded into 96-well plates (100 μL of 200,000 cells/mL/well). Cells were immediately stimulated with PMA and treated with 0.5 mg/mL A1AT or 1/10 MSCs or their combination. The fluorescent and phase contrast images were taken by the FV3000 confocal laser scanning microscope (Olympus).

### PBMC flow cytometry assay

Pooled human PBMCs were purchased from Zenbio and recovered overnight before stimulation. LPS (100 ng/mL) and 25 uL human CD3/CD28 activator solution / million cells and the treatments were added for 72 hours. Then PBMCs were cultured with 1x Cell Stimulation Cocktail plus protein transport inhibitors (Invitrogen) for 4 hrs. Single cells were harvested and stained with anti-human CD3-APCcy7 and CD14-FITC for 15 mins at room temperature. After that, the cells were fixed and permeabilized with the BD Cytofix/Cytoperm™ Fixation/Permeabilization Solution Kit (BD Bioscience) and labeled intracellularly with anti-human IFNγ-APC, TNFα-BV605 (Biolegend) and IL10-PE (ebioscience). Data were collected on Attune NxT Flow Cytometer (Thermofisher) and analyzed using FlowJo software.

### Phagocytosis analysis

FITC-labeled pHrodo E. coli Bioparticles^®^ Conjugate (Thermo Fisher) were used to assess phagocytosis of THP1-derived macrophage and HL-60 derived neutrophils. The stimulation and treatment methods were described in their inflammation assay paragraph.

E. coli particles were resuspended in PBS and coated with rabbit polyclonal IgG antibodies (Escherichia coli BioParticles™ Opsonizing Reagent, Thermo Fisher) at 37°C for 1 hr. Next, cells were incubated with 0.1 mg/mL coated E. coli particles at 37°C for 3 hrs. Non-phagocytosed E. coli bioparticles were removed by washing with PBS (PH=7.4). Next, cells were fixed with 4% PFA, permeabilized with 0.05% TritonX-100, and stained in DAPI solution. Cells were imaged with Olympus FV3000 confocal microscope and analyzed using ImageJ software.

### Acute lung injury and inflammation mice

All animal experiments were approved by the Animal Care and Use Committee of the University of Nebraska-Lincoln. 10-week old male C57BL/6 mice (25 g) were purchased from Jackson Lab. For A1AT treatment, 2 mg A1AT (in 200 μL PBS) was injected intraperitoneally (i.p.) at 48 hrs, 24 hrs, and 0 hr before the LPS challenge (three doses). Mice were anesthetized with ketamine (120 mg/kg body weight or BW, i.p.) and xylazine (16 mg/kg BW, i.p.). Mice were placed in the prone position. A 22 gauge (G) venous catheter was gently inserted into the trachea along the tongue’s root in the vertical direction. Approximately 10 mm of the catheter was inserted. 50 μL of LPS was instilled. For survival rate assay, 20 mg LPS/kg BW was used. For lung tissue injury and cytokine production studies, 10 mg LPS/kg BW was used. Using a pipette, 1×10^6^ MSCs were instilled via the catheter 30 mins after the LPS challenge. Next, 1 mL air was instilled to ensure LPS and cells were distributed well in the lung. The mouse’s upper body was kept upright for 30 seconds to avoid fluid leakage. The body temperature was maintained at 37°C until full awareness. The mouse was transferred to ventilated cage individually with free access to food and water. The survival rate and body weight were monitored and recorded twice a day.

### Bronchoalveolar lavage fluid (BALF) and tissue harvest

Anesthesia was induced. The trachea was carefully exposed, and a 22 G venous catheter was inserted after a 5 mm cut to the trachea. 0.5 mL PBS was instilled, followed by 0.1 mL of air. After 60s, the fluid was aspirated. This process was repeated three times to collect all BALF. Cells in BALF were harvested by centrifuging at 300 g for 10 mins. BALF cells were resuspended using 90% FBS plus 10% DMSO and frozen in a Mr. Frost at - 80°C before long-term storage in liquid nitrogen. The supernatant was frozen at −80°C for cytokine analysis. After collecting BALF, lungs and other organs were harvested and fixed in 4% PFA for histology analyses.

### Histology and immune staining

The fixed tissues were embedded in paraffin and sectioned (5 μm thickness). Sections were dewaxed with the Leica Auto Stainer XL and soaked in EDTA pH 8.0 (Abcam) or 10 mM Sodium Citrate solution pH 6.0 (Invitrogen) for antigen retrieval. The TBS superblock blocking buffer(Thermo Fisher) was applied to the slide for 1 hr, followed by primary antibody incubation overnight at 4 °C. Slides were washed with PBS and incubated with secondary antibody and DAPI at room temperature in the dark.

### BALF cells staining

Cells collected from BALF were thawed, resuspended in PBS, and fixed in 4% PFA for 20 mins. Next, cells were washed in dd H2O, placed on a Poly-Prep Slide (Sigma), and heated until dry. Slides were blocked and stained as the tissue immune staining.

### TUNEL staining

The One-step TUNEL In Situ Apoptosis AF 594 Kit (Elabscience) was used. Paraffin sections were dewaxed and treated with 1x proteinase K solution at 37°C for 20 mins. Next, sections were labeled by TDT reaction mixture for 2 hrs at 37°C. The reaction was stopped with PBS and stained with DAPI before mounting and imaging.

### Statistical analysis

All the data were analyzed using GraphPad Prism 8 statistical software and shown as mean ± standard error of the mean. P value was determined by one-way analysis of variance (ANOVA) for comparison between the means of three or more groups, log-rank test for survival, or unpaired two-tailed t-tests for two groups analysis. The significance levels are indicated by p-value, *: p<0.05, **: p<0.01, ***: p<0.001.

## Author contribution

Conceptualization: YL, WV, CD, YW, MA, EH, AS, YL, RJ, GL; Investigation: LH, XW, OW; Data analysis: LH, XW, XL. Writing-original draft: YL, LH. Writing-review and editing: LH, EH, AS, YL

## Completing interest

Dr. Lei owns equity in CellGro Technologies, LLC. This financial interest has been reviewed by the University’s Individual Conflict of Interest Committee and is currently being managed by the University.

## REFERENCES

1. Fajgenbaum, D. C. & June, C. H. Cytokine Storm. N. Engl. J. Med. 383, 2255–2273 (2020).

2. Hussen, J., Kandeel, M., Hemida, M. G. & Al-Mubarak, A. I. A. Antibody-based immunotherapeutic strategies for COVID-19. Pathogens 9, 1–18 (2020).

3. Kiselevskiy, M. et al. Immune pathogenesis of covid-19 intoxication: Storm or silence? Pharmaceuticals 13, 1–17 (2020).

4. Nouveau, L. et al. Immunological analysis of the murine anti-CD3-induced cytokine release syndrome model and therapeutic efficacy of anti-cytokine antibodies. Eur. J. Immunol. 51, 2074–2085 (2021).

5. Kash, J. C. et al. Genomic analysis of increased host immune and cell death responses induced by 1918 influenza virus. Nature 443, 578–581 (2006).

6. Morgan, R. A. et al. Case report of a serious adverse event following the administration of t cells transduced with a chimeric antigen receptor recognizing ERBB2. Mol. Ther. 18, 843–851 (2010).

7. A current view on inflammation. Nat. Immunol. 18, 825 (2017).

8. Netea, M. G. et al. A guiding map for inflammation. Nat. Immunol. 18, 826–831 (2017).

9. Fullerton, J. N. & Gilroy, D. W. Resolution of inflammation: A new therapeutic frontier. Nat. Rev. Drug Discov. 15, 551–567 (2016).

10. Feehan, K. T. & Gilroy, D. W. Is Resolution the End of Inflammation? Trends Mol. Med. 25, 198–214 (2019).

11. Mahmudpour, M., Roozbeh, J., Keshavarz, M., Farrokhi, S. & Nabipour, I. COVID-19 cytokine storm: The anger of inflammation. Cytokine 133, 155151 (2020).

12. Ghanbarpour, R. et al. Pulmonary infections in ICU patients without underlying disease on ventilators. Trauma Mon. 19, 41–44 (2014).

13. Motwani, M. P. et al. Potent Anti-Inflammatory and Pro-Resolving Effects of Anabasum in a Human Model of Self-Resolving Acute Inflammation. Clin. Pharmacol. Ther. 104, 675–686 (2018).

14. Zhang, Z. et al. Mesenchymal stem cells promote the resolution of cardiac inflammation after ischemia reperfusion via enhancing efferocytosis of neutrophils. J. Am. Heart Assoc. 9, (2020).

15. Giugliano, G. R., Giugliano, R. P., Gibson, C. M. & Kuntz, R. E. Meta-analysis of corticosteroid treatment in acute myocardial infarction. Am. J. Cardiol. 91, 1055–1059 (2003).

16. Proto, J. D. et al. Regulatory T Cells Promote Macrophage Efferocytosis during Inflammation Resolution. Immunity 49, 666–677.e6 (2018).

17. Mietto, B. S. et al. Role of IL-10 in resolution of inflammation and functional recovery after peripheral nerve injury. J. Neurosci. 35, 16431–16442 (2015).

18. Hutchins, A. P., Diez, D. & Miranda-Saavedra, D. The IL-10/STAT3-mediated anti-inflammatory response: Recent developments and future challenges. Brief. Funct. Genomics 12, 489–498 (2013).

19. Gu, Z. et al. Resolvin D1, resolvin D2 and maresin 1 activate the GSK3β anti-inflammatory axis in TLR4-engaged human monocytes. Innate Immun. 22, 186–195 (2016).

20. Cioccari, L., Luethi, N. & Masoodi, M. Lipid Mediators in Critically Ill Patients: A Step Towards Precision Medicine. Front. Immunol. 11, 1–10 (2020).

21. Serhan, C. N., Chiang, N., Dalli, J. & Levy, B. D. Lipid mediators in the resolution of inflammation. Cold Spring Harb. Perspect. Biol. 7, 1–20 (2015).

22. Fonseca, M. T. et al. A leukotriene-dependent spleen-liver axis drives TNF production in systemic inflammation. Sci. Signal. 14, (2021).

23. Gupta, N. et al. Mesenchymal stem cells enhance survival and bacterial clearance in murine Escherichia coli pneumonia. Thorax 67, 533–539 (2012).

24. Németh, K. et al. Bone marrow stromal cells attenuate sepsis via prostaglandin E 2-dependent reprogramming of host macrophages to increase their interleukin-10 production. Nat. Med. 15, 42–49 (2009).

25. Pedrazza, L. et al. Mesenchymal stem cells improves survival in LPS-induced acute lung injury acting through inhibition of NETs formation. J. Cell. Physiol. 232, 3552–3564 (2017).

26. Perlee, D. et al. Human Adipose-Derived Mesenchymal Stem Cells Modify Lung Immunity and Improve Antibacterial Defense in Pneumosepsis Caused by Klebsiella pneumoniae. Stem Cells Transl. Med. 8, 785–796 (2019).

27. Shin, S. et al. The therapeutic effect of human adult stem cells derived from adipose tissue in endotoxemic rat model. Int. J. Med. Sci. 10, 8–18 (2012).

28. Devaney, J. et al. Human mesenchymal stromal cells decrease the severity of acute lung injury induced by E. Coli in the rat. Thorax 70, 625–635 (2015).

29. Yang, Y. et al. The Vascular Endothelial Growth Factors-Expressing Character of Mesenchymal Stem Cells Plays a Positive Role in Treatment of Acute Lung Injury In Vivo. Mediators Inflamm. 2016, (2016).

30. For, E. Treatment With Human Wharton ‘ s Jelly-Derived Mesenchymal Stem Cells Attenuates Sepsis-Induced Kidney Injury, Liver Injury, and E. 1048–1057 (2016).

31. Danchuk, S. et al. Human multipotent stromal cells attenuate lipopolysaccharide-induced acute lung injury in mice via secretion of tumor necrosis factor-α-induced protein 6. Stem Cell Res. Ther. 2, 1–15 (2011).

32. Curley, G. F. et al. Mesenchymal stem cells enhance recovery and repair following ventilator-induced lung injury in the rat. Thorax 67, 496–501 (2012).

33. Hackstein, H. et al. Prospectively defined murine mesenchymal stem cells inhibit Klebsiella pneumoniae-induced acute lung injury and improve pneumonia survival. Respir. Res. 16, 1–12 (2015).

34. Alcayaga-Miranda, F. et al. Combination therapy of menstrual derived mesenchymal stem cells and antibiotics ameliorates survival in sepsis. Stem Cell Res. Ther. 6, 1–13 (2015).

35. Curley, G. F. et al. Effects of intratracheal mesenchymal stromal cell therapy during recovery and resolution after ventilator-induced lung injury. Anesthesiology 118, 924–933 (2013).

36. Hayes, M. et al. Therapeutic efficacy of human mesenchymal stromal cells in the repair of established ventilator-induced lung injury in the rat. Anesthesiology 122, 363–373 (2015).

37. Rocheteau, P. et al. sepsis induces long-term metabolic and mitochondrial muscle stem cell dysfunction amenable by mesenchymal stem cell therapy. Nat. Commun. 6, 1–12 (2015).

38. Jackson, M. V. et al. Mitochondrial Transfer via Tunneling Nanotubes is an Important Mechanism by Which Mesenchymal Stem Cells Enhance Macrophage Phagocytosis in the In Vitro and In Vivo Models of ARDS. Stem Cells 34, 2210–2223 (2016).

39. Krasnodembskaya, A. et al. Human mesenchymal stem cells reduce mortality and bacteremia in gram-negative sepsis in mice in part by enhancing the phagocytic activity of blood monocytes. Am. J. Physiol. - Lung Cell. Mol. Physiol. 302, 1003–1013 (2012).

40. Morrison, T. J. et al. Mesenchymal stromal cells modulate macrophages in clinically relevant lung injury models by extracellular vesicle mitochondrial transfer. Am. J. Respir. Crit. Care Med. 196, 1275–1286 (2017).

41. Li, B. et al. Bone marrow mesenchymal stem cells protect alveolar macrophages from lipopolysaccharide-induced apoptosis partially by inhibiting the Wnt/β-catenin pathway. Cell Biol. Int. 39, 192–200 (2015).

42. Lee, J. W. et al. Therapeutic effects of human mesenchymal stem cells in ex vivo human lungs injured with live bacteria. Am. J. Respir. Crit. Care Med. 187, 751–760 (2013).

43. Hall, S. R. R. et al. Mesenchymal stromal cells improve survival during sepsis in the absence of heme oxygenase-1: The importance of neutrophils. Stem Cells 31, 397–407 (2013).

44. Laroye, C. et al. Bone marrow vs Wharton’s jelly mesenchymal stem cells in experimental sepsis: A comparative study. Stem Cell Res. Ther. 10, 1–11 (2019).

45. Jerkic, M. et al. Human Umbilical Cord Mesenchymal Stromal Cells Attenuate Systemic Sepsis in Part by Enhancing Peritoneal Macrophage Bacterial Killing via Heme Oxygenase-1 Induction in Rats. Anesthesiology 132, 140–154 (2020).

46. Rabani, R. et al. Mesenchymal stem cells enhance NOX2-dependent reactive oxygen species production and bacterial killing in macrophages during sepsis. Eur. Respir. J. 51, 1–14 (2018).

47. Kim, J. & Hematti, P. Mesenchymal stem cell-educated macrophages: A novel type of alternatively activated macrophages. Exp. Hematol. 37, 1445–1453 (2009).

48. Mao, Y. X. et al. Adipose tissue-derived mesenchymal stem cells attenuate pulmonary infection caused by Pseudomonas aeruginosa via inhibiting overproduction of prostaglandin E2. Stem Cells 33, 2331–2342 (2015).

49. Mei, S. H. J. et al. Mesenchymal stem cells reduce inflammation while enhancing bacterial clearance and improving survival in sepsis. Am. J. Respir. Crit. Care Med. 182, 1047–1057 (2010).

50. Sung, D. K. et al. Antibacterial effect of mesenchymal stem cells against Escherichia coli is mediated by secretion of beta-defensin-2 via toll-like receptor 4 signalling. Cell. Microbiol. 18, 424–436 (2016).

51. Lu, Z. et al. Mesenchymal stem cells induce dendritic cell immune tolerance via paracrine hepatocyte growth factor to alleviate acute lung injury. Stem Cell Res. Ther. 10, 1–16 (2019).

52. Silva, J. D. et al. Eicosapentaenoic acid potentiates the therapeutic effects of adipose tissue-derived mesenchymal stromal cells on lung and distal organ injury in experimental sepsis. Stem Cell Res. Ther. 10, 1–16 (2019).

53. Marrow, B. et al. induced Acute Lung Injury and Enhance Resolution of Ventilator-induced Lung Injury in Rats. 502–516 (2018).

54. Zhang, Z. et al. Combination therapy of human umbilical cord mesenchymal stem cells and FTY720 attenuates acute lung injury induced by lipopolysaccharide in a murine model. Oncotarget 8, 77407–77414 (2017).

55. Pati, S. et al. Bone marrow derived mesenchymal stem cells inhibit inflammation and preserve vascular endothelial integrity in the lungs after hemorrhagic shock. PLoS One 6, (2011).

56. Asmussen, S. et al. Human mesenchymal stem cells reduce the severity of acute lung injury in a sheep model of bacterial pneumonia. Thorax 69, 819–825 (2014).

57. Thompson, M. et al. Cell therapy with intravascular administration of mesenchymal stromal cells continues to appear safe: An updated systematic review and meta-analysis. EClinicalMedicine 19, (2020).

58. Le Blanc, K. et al. Mesenchymal stem cells for treatment of steroid-resistant, severe, acute graft-versus-host disease: a phase II study. Lancet 371, 1579–1586 (2008).

59. Karussis, D. et al. Safety and immunological effects of mesenchymal stem cell transplantation in patients with multiple sclerosis and amyotrophic lateral sclerosis. Arch. Neurol. 67, 1187–1194 (2010).

60. He, X. et al. Umbilical cord-derived mesenchymal stem (stromal) cells for treatment of severe sepsis: aphase 1 clinical trial. Transl. Res. 199, 52–61 (2018).

61. McIntyre, L. A. et al. Cellular immunotherapy for septic shock: A phase I clinical trial. Am. J. Respir. Crit. Care Med. 197, 337–347 (2018).

62. Chen, J. et al. Clinical Study of Mesenchymal Stem Cell Treatment for Acute Respiratory Distress Syndrome Induced by Epidemic Influenza A (H7N9) Infection: A Hint for COVID-19 Treatment. Engineering 6, 1153–1161 (2020).

63. Connick, P. et al. Autologous mesenchymal stem cells for the treatment of secondary progressive multiple sclerosis: An open-label phase 2a proof-of-concept study. Lancet Neurol. 11, 150–156 (2012).

64. Wilson, J. G. et al. Mesenchymal stem (stromal) cells for treatment of ARDS: A phase 1 clinical trial. Lancet Respir. Med. 3, 24–32 (2015).

65. Panés, J. et al. Expanded allogeneic adipose-derived mesenchymal stem cells (Cx601) for complex perianal fistulas in Crohn’s disease: a phase 3 randomised, double-blind controlled trial. Lancet 388, 1281–1290 (2016).

66. Kebriaei, P. et al. A Phase 3 Randomized Study of Remestemcel-L versus Placebo Added to Second-Line Therapy in Patients with Steroid-Refractory Acute Graft-versus-Host Disease. Biol. Blood Marrow Transplant. 26, 835–844 (2020).

67. Zheng, G. et al. Treatment of acute respiratory distress syndrome with allogeneic adipose-derived mesenchymal stem cells: A randomized, placebo-controlled pilot study. Respir. Res. 15, 1–10 (2014).

68. Lv, H. et al. Mesenchymal stromal cells as a salvage treatment for confirmed acute respiratory distress syndrome: preliminary data from a single-arm study. Intensive Care Med. 46, 1944–1947 (2020).

69. Matthay, M. A. et al. Treatment with allogeneic mesenchymal stromal cells for moderate to severe acute respiratory distress syndrome (START study): a randomised phase 2a safety trial. Lancet Respir. Med. 7, 154–162 (2019).

70. Gennadiy, G. et al. The Results of the Single Center Pilot Randomized Russian Clinical Trial of Mesenchymal Stromal Cells in Severe Neutropenic Patients with Septic Shock (RUMCESS). Int. J. Blood Res. Disord. 5, (2018).

71. Rossello-Gelabert, M., Gonzalez-Pujana, A., Igartua, M., Santos-Vizcaino, E. & Hernandez, R. M. Clinical progress in MSC-based therapies for the management of severe COVID-19. Cytokine Growth Factor Rev. 68, 25–36 (2022).

72. Liang, B. et al. Clinical remission of a critically ill COVID-19 patient treated by human umbilical cord mesenchymal stem cells. ChinaXiv (2020) doi:10.3969/j.issn.2095-4344.2012.49.011.

73. Leng, Z. et al. Transplantation of ACE2-Mesenchymal stem cells improves the outcome of patients with covid-19 pneumonia. Aging Dis. 11, 216–228 (2020).

74. Lanzoni, G. et al. Umbilical cord mesenchymal stem cells for COVID-19 acute respiratory distress syndrome: A double-blind, phase 1/2a, randomized controlled trial. Stem Cells Transl. Med. 10, 660–673 (2021).

75. Sánchez-Guijo, F. et al. Adipose-derived mesenchymal stromal cells for the treatment of patients with severe SARS-CoV-2 pneumonia requiring mechanical ventilation. A proof of concept study. EClinicalMedicine 25, (2020).

76. Chen, X., Shan, Y., Wen, Y., Sun, J. & Du, H. Mesenchymal stem cell therapy in severe COVID-19: A retrospective study of short-term treatment efficacy and side effects. Journal of Infection vol. 81 (2020).

77. Gorman, E., Millar, J., McAuley, D. & O’Kane, C. Mesenchymal stromal cells for acute respiratory distress syndrome (ARDS), sepsis, and COVID-19 infection: optimizing the therapeutic potential. Expert Rev. Respir. Med. 15, 301–324 (2021).

78. Guttman, O., S Freixo-Lima, G. & C Lewis, E. Alpha1-antitrypsin, an endogenous immunoregulatory molecule: distinction between local and systemic effects on tumor immunology. Integr. Cancer Sci. Ther. 2, 272–280 (2016).

79. Bergin, D. A., Hurley, K., McElvaney, N. G. & Reeves, E. P. Alpha-1 antitrypsin: A potent anti-inflammatory and potential novel therapeutic agent. Arch. Immunol. Ther. Exp. (Warsz). 60, 81–97 (2012).

80. Toldo, S. et al. Alpha-1 antitrypsin inhibits caspase-1 and protects from acute myocardial ischemia-reperfusion injury. J. Mol. Cell. Cardiol. 51, 244–251 (2011).

81. Janciauskiene, S. et al. The Multifaceted Effects of Alpha1-Antitrypsin on Neutrophil Functions. Front Pharmacol. 17, 341 (2018).

82. Marcondes, A. M. et al. Inhibition of IL-32 activation by α-1 antitrypsin suppresses alloreactivity and increases survival in an allogeneic murine marrow transplantation model. Blood. 118, 5031–9 (2011).

83. Shapiro, S. D. et al. Neutrophil Elastase Contributes to Cigarette Smoke-Induced Emphysema in Mice. Am. J. Pathol. 163, 2329–2335 (2003).

84. Kidokoro Y, Kravis TC, Moser KM, Taylor JC, C. I. Relationship of Leukocyte Elastase Concentration to Severity of Emphysema in Homozygous α1-Antitrypsin-Deficient Persons. Am Rev Respir Dis. 115, 793–803 (1977).

85. Yuan-Ping Han, Chunli Yan, and W. L. G. Proteolytic Activation of Matrix Metalloproteinase-9 in Skin Wound Healing Is Inhibited by α-1-Antichymotrypsin. J Invest Dermatol. 128, 2334–2342 (2008).

86. He, S., Chen, H. & Zheng, J. Inhibition of tryptase and chymase induced nucleated cell infiltration by proteinase inhibitors 1. Acta Pharmacol Sin. 25, 1677–1684 (2004).

87. Bergin, D. A. et al. α-1 antitrypsin regulates human neutrophil chemotaxis induced by soluble immune complexes and IL-8. J. Clin. Invest. 120, 4236–4250 (2010).

88. Jedicke, N. et al. α-1-antitrypsin inhibits acute liver failure in mice. Hepatology. 59, 2299–2308 (2014).

89. Libert, C., Molle, W. Van, Brouckaert, P. & Fiers, W. Alpha-1-Antitrypsin Inhibits the Lethal Response to TNF in Mice. J. Immunol. 157, 5126–5129 (1996).

90. Subramaniyam, D. et al. Effects of alpha 1-antitrypsin on endotoxin-induced lung inflammation in vivo. Inflamm. Res. 59, 571–578 (2010).

91. Griese, M. et al. α1-Antitrypsin inhalation reduces airway inflammation in cystic fibrosis patients. Eur. Respir. J. 29, 240–250 (2007).

92. Pott, G. B., Chan, E. D., Dinarello, C. A. & Shapiro, L. α-1-Antitrypsin is an endogenous inhibitor of pro-inflammatory cytokine production in whole blood. J. Leukoc. Biol. 85, 886–895 (2009).

93. Ochayon, D. E., Mizrahi, M., Shahaf, G., Baranovski, B. M. & Lewis, E. C. Human α1-Antitrypsin Binds to Heat-Shock Protein gp96 and Protects from Endogenous gp96-Mediated Injury In vivo. Front. Immunol. 4, 320 (2013).

94. Tilg, B. H., Vannier, E., Vachino, G., Dinardlo, C. A. & Mier, J. W. Anti-inflammatory properties of hepatic acute phase proteins: preferential induction of interleukin 1 (IL-1) receptor antagonist over IL-1 beta synthesis by human peripheral blood mononuclear cells. J Exp Med. 178, 1629–36 (1993).

95. Finotti, P. & Pagetta, A. A heat shock protein70 fusion protein with alpha1-antitrypsin in plasma of type 1 diabetic subjects. Biochem Biophys Res Commun. 315, 297–305 (2004).

96. Lockett, A. D. et al. α_1_-Antitrypsin modulates lung endothelial cell inflammatory responses to TNF-α. Am J Respir Cell Mol Biol. 49, 143–50 (2013).

97. Chan, E. D. et al. Alpha-1-antitrypsin inhibits nitric oxide production. J. Leukoc. Biol. 92, 1251–1260 (2012).

98. Zhou, T. et al. Alpha-1 antitrypsin attenuates M1 microglia-mediated neuroinflammation in retinal degeneration. Front. Immunol. 9, 1202 (2018).

99. Jonigk, D. et al. Anti-inflammatory and immunomodulatory properties of 1-antitrypsin without inhibition of elastase. Proc. Natl. Acad. Sci. 110, 15007–15012 (2013).

100. Serban, K. A. et al. Alpha-1 antitrypsin supplementation improves alveolar macrophages efferocytosis and phagocytosis following cigarette smoke exposure. PLoS One 12, 1–17 (2017).

101. Nita, I. M., Serapinas, D. & Janciauskiene, S. M. α1-Antitrypsin regulates CD14 expression and soluble CD14 levels in human monocytes in vitro. Int. J. Biochem. Cell Biol. 39, 1165–1176 (2007).

102. Janciauskiene, S. M., Nita, I. M. & Stevens, T. α1-antitrypsin, old dog, new tricks: α1-antitrypsin exerts in vitro anti-inflammatory activity in human monocytes by elevating cAMP. J. Biol. Chem. 282, 8573–8582 (2007).

103. Ozeri, E., Mizrahi, M., Shahaf, G. & Lewis, E. C. -1 Antitrypsin Promotes Semimature, IL-10-Producing and Readily Migrating Tolerogenic Dendritic Cells. J. Immunol. 189, 146–153 (2012).

104. Churg, A. et al. Alpha-1-Antitrypsin and a Broad Spectrum Acute Anti-inflammatory Effects. Lab Invest 81, 1119–1131 (2001).

105. Kaner, Z. et al. Acute Phase Protein α1-Antitrypsin Reduces the Bacterial Burden in Mice by Selective Modulation of Innate Cell Responses. J. Infect. Dis. 211, 1489–1498 (2015).

106. Pott, G. B., Beard, K. S., Bryan, C. L., Merrick, D. T. & Shapiro, L. Alpha-1 Antitrypsin Reduces Severity of Pseudomonas Pneumonia in Mice and Inhibits Epithelial Barrier Disruption and Pseudomonas Invasion of Respiratory Epithelial Cells. Front. Public Heal. 1, 1–13 (2013).

107. Wanner, A., Arce, A. De & Pardee, E. Novel therapeutic uses of alpha-1 antitrypsin: A window to the future. COPD J. Chronic Obstr. Pulm. Dis. 9, 583–588 (2012).

108. Jia, Q. et al. Short cyclic peptides derived from the C-terminal sequence of α1-antitrypsin exhibit significant anti-HIV-1 activity. Bioorganic Med. Chem. Lett. 22, 2393–2395 (2012).

109. Bristow CL, Modarresi R, Babayeva MA, LaBrunda M, Mukhtarzad R, Trucy M, Franklin A, Reeves RE, Long A, Mullen MP, Cortes J, W. R. A feedback regulatory pathway between LDL and alpha-1 proteinase inhibitor in chronic inflammation and infection. Discov Med. 16, 201–18 (2013).

110. Bristow, C. L., Babayeva, M. A., LaBrunda, M., Mullen, M. P. & Winston, R. α 1proteinase inhibitor regulates CD4 + lymphocyte levels and is rate limiting in HIV-1 disease. PLoS One 7, 1–10 (2012).

111. Abdulsalam, S. I., Abdulatif, A., Joyal, J., Wisam, G. & Ajayeb, A. Increased Prevalence of the Alpha-1-Antitrypsin (A1AT) Deficiency-Related S Gene in Patients Infected With Human Immunodeficiency Virus Type 1. J. Med. Virol. 81, 1047–1051 (2009).

112. Münch, J. et al. Discovery and Optimization of a Natural HIV-1 Entry Inhibitor Targeting the gp41 Fusion Peptide. Cell 129, 263–275 (2007).

113. Shapiro, L., Pott, G. B. & Ralston, A. H. Alpha-1-antitrypsin inhibits human immunodeficiency virus type 1. FASEB J. 15, 115–122 (2002).

114. Moldthan, H. L. et al. Alpha 1-antitrypsin therapy mitigated ischemic stroke damage in rats. J. Stroke Cerebrovasc. Dis. 23, e355–e363 (2014).

115. Koulmanda, M. et al. Alpha 1-antitrypsin reduces inflammation and enhances mouse pancreatic islet transplant survival. Proc. Natl. Acad. Sci. 109, 15443–15448 (2012).

116. Petrache, I. et al. Alpha-1 Antitrypsin Inhibits Antitrypsin Inhibits Caspase-3 Activity, Preventing Lung Endothelial Cell Apoptosis. Am. J. Pathol. 169, 1155–1166 (2006).

117. Kalis, M., Kumar, R., Janciauskiene, S., Salehi, A. & Cilio, C. M. A 1-Antitrypsin Enhances Insulin Secretion and Prevents Cytokine-Mediated Apoptosis in Pancreatic B-Cells. Islets 2, 185–189 (2010).

118. Bellacen, K., Kalay, N., Ozeri, E., Shahaf, G. & Lewis, E. C. Revascularization of pancreatic islet allografts is enhanced by α-1-Antitrypsin under anti-inflammatory conditions. Cell Transplant. 22, 2119–2133 (2013).

119. Aldonyte, R. et al. Endothelial alpha-1-antitrypsin attenuates cigarette smoke induced apoptosis in vitro. COPD J. Chronic Obstr. Pulm. Dis. 5, 153–162 (2008).

120. Janciauskiene, S. & Welte, T. Well-known and less well-known functions of Alpha-1 antitrypsin: Its role in chronic obstructive pulmonary disease and other disease developments. Ann. Am. Thorac. Soc. 13, S280--S288 (2016).

121. Kim, M., Cai, Q. & Oh, Y. Therapeutic potential of alpha-1 antitrypsin in human disease. Ann. Pediatr. Endocrinol. Metab. 23, 131–135 (2018).

122. Ritzmann, F. et al. Therapeutic application of alpha-1 antitrypsin in COVID-19. Am. J. Respir. Crit. Care Med. 204, 224–227 (2021).

123. Philippe, A. et al. Imbalance between alpha-1-antitrypsin and interleukin 6 is associated with in-hospital mortality and thrombosis during COVID-19. Biochimie 202, (2022).

124. Bai, X. et al. Hypothesis: Alpha-1-antitrypsin is a promising treatment option for COVID-19. Med. Hypotheses 146, 110394 (2021).

125. McEvoy, N. L. et al. A randomised, double-blind, placebo-controlled, pilot trial of intravenous plasma purified alpha-1 antitrypsin for SARS-CoV-2-induced Acute Respiratory Distress Syndrome: a structured summary of a study protocol for a randomised, controlled trial. Trials 22, 22–24 (2021).

126. McElvaney, O. J. et al. A randomized, double-blind, placebo-controlled trial of intravenous alpha-1 antitrypsin for ARDS secondary to COVID-19. Med 3, 233–248.e6 (2022).

127. Schuster, R. et al. Distinct anti-inflammatory properties of alpha1-antitrypsin and corticosteroids reveal unique underlying mechanisms of action. Cell. Immunol. 356, 104177 (2020).

128. Jiang, D. et al. Suppression of Neutrophil-Mediated Tissue Damage—A Novel Skill of Mesenchymal Stem Cells. Stem Cells 34, 2393–2406 (2016).

129. Kim, E. Y. et al. Post-sepsis immunosuppression depends on NKT cell regulation of mTOR/IFN-γ in NK cells. J. Clin. Invest. 130, 3238–3252 (2020).

130. Hortová-Kohoutková, M. et al. Phagocytosis-Inflammation Crosstalk in Sepsis: New Avenues for Therapeutic Intervention. Shock 54, 606–614 (2020).

131. Jin, Z. et al. TRIM59 Protects Mice From Sepsis by Regulating Inflammation and Phagocytosis in Macrophages. Front. Immunol. 11, 1–12 (2020).

132. de Witte, S. F. H. et al. Immunomodulation By Therapeutic Mesenchymal Stromal Cells (MSC) Is Triggered Through Phagocytosis of MSC By Monocytic Cells. Stem Cells 36, 602–615 (2018).

133. Yip, H. K. et al. Human Umbilical Cord-Derived Mesenchymal Stem Cells for Acute Respiratory Distress Syndrome. Crit. Care Med. E391–E399 (2020) doi: 10.1097/CCM.0000000000004285.

134. Barkama, R. et al. Placenta-Derived Cell Therapy to Treat Patients With Respiratory Failure Due to Coronavirus Disease 2019. Crit. Care Explor. 2, e0207 (2020).

135. Zhu, R. et al. Mesenchymal stem cell treatment improves outcome of COVID-19 patients via multiple immunomodulatory mechanisms. Cell Res. 31, 1244–1262 (2021).

136. Tang, L. et al. Clinical study using mesenchymal stem cells for the treatment of patients with severe COVID-19. Front. Med. 14, 664–673 (2020).

137. Shu, L. et al. Treatment of severe COVID-19 with human umbilical cord mesenchymal stem cells. Stem Cell Res. Ther. 11, 1–11 (2020).

138. Adas, G. et al. The Systematic Effect of Mesenchymal Stem Cell Therapy in Critical COVID-19 Patients: A Prospective Double Controlled Trial. Cell Transplant. 30, 1–14 (2021).

139. Xu, X. et al. Evaluation of the safety and efficacy of using human menstrual blood-derived mesenchymal stromal cells in treating severe and critically ill COVID-19 patients: An exploratory clinical trial. Clin. Transl. Med. 11, (2021).

140. Meng, F. et al. Human umbilical cord-derived mesenchymal stem cell therapy in patients with COVID-19: a phase 1 clinical trial. Signal Transduct. Target. Ther. 5, (2020).

141. Kavianpour, M., Saleh, M. & Verdi, J. The role of mesenchymal stromal cells in immune modulation of COVID-19: Focus on cytokine storm. Stem Cell Res. Ther. 11, (2020).

142. Grom, A. A., Horne, A. & De Benedetti, F. Macrophage activation syndrome in the era of biologic therapy. Nat. Rev. Rheumatol. 12, 259–268 (2016).

143. Grigorieva, K. N. et al. Macrophage activation syndrome in COVID-19. Obstet. Gynecol. Reprod. 15, 313–320 (2021).

144. Crayne, C. B., Albeituni, S., Nichols, K. E. & Cron, R. Q. The immunology of macrophage activation syndrome. Front. Immunol. 10, 1–11 (2019).

145. McGonagle, D., Ramanan, A. V. & Bridgewood, C. Immune cartography of macrophage activation syndrome in the COVID-19 era. Nat. Rev. Rheumatol. 17, 145–157 (2021).

146. Ackermann, M. et al. Patients with COVID-19: in the dark-NETs of neutrophils. Cell Death Differ. 28, 3125–3139 (2021).

147. Chiang, C. C., Korinek, M., Cheng, W. J. & Hwang, T. L. Targeting Neutrophils to Treat Acute Respiratory Distress Syndrome in Coronavirus Disease. Front. Pharmacol. 11, (2020).

148. Skendros, P. et al. Complement and tissue factor–enriched neutrophil extracellular traps are key drivers in COVID-19 immunothrombosis. J. Clin. Invest. 130, 6151–6157 (2020).

149. McKenna, E. et al. Neutrophils in COVID-19: Not Innocent Bystanders. Front. Immunol. 13, 1–12 (2022).

150. Reusch, N. et al. Neutrophils in COVID-19. Front. Immunol. 12, 1–9 (2021).

151. Meizlish, M. L. et al. A neutrophil activation signature predicts critical illness and mortality in COVID-19. Blood Adv. 5, 1164–1177 (2021).

152. Masso-Silva, J. A. et al. Increased Peripheral Blood Neutrophil Activation Phenotypes and Neutrophil Extracellular Trap Formation in Critically Ill Coronavirus Disease 2019 (COVID-19) Patients: A Case Series and Review of the Literature. Clin. Infect. Dis. 74, 479–489 (2022).

153. Dowey, R. et al. Enhanced neutrophil extracellular trap formation in COVID-19 is inhibited by the protein kinase C inhibitor ruboxistaurin. ERJ Open Res. 8, (2022).

154. Narasaraju, T. et al. Neutrophilia and NETopathy as Key Pathologic Drivers of Progressive Lung Impairment in Patients With COVID-19. Front. Pharmacol. 11, 1–8 (2020).

155. Laforge, M. et al. Tissue damage from neutrophil-induced oxidative stress in COVID-19. Nat. Rev. Immunol. 20, 515–516 (2020).

156. Al-Kuraishy, H. M. et al. Neutrophil Extracellular Traps (NETs) and Covid-19: A new frontiers for therapeutic modality. Int. Immunopharmacol. 104, (2022).

157. Yaqinuddin, A., Kvietys, P. & Kashir, J. Since January 2020 Elsevier has created a COVID-19 resource centre with free information in English and Mandarin on the novel coronavirus COVID-19. The COVID-19 resource centre is hosted on Elsevier Connect, the company ‘ s public news and information. (2020).

158. Lefrançais, E., Mallavia, B., Zhuo, H., Calfee, C. S. & Looney, M. R. Maladaptive role of neutrophil extracellular traps in pathogen-induced lung injury. JCI insight 3, 1–15 (2018).

159. Yaqinuddin, A. & Kashir, J. Novel therapeutic targets for SARS-CoV-2-induced acute lung injury: Targeting a potential IL-1β/neutrophil extracellular traps feedback loop. Med. Hypotheses 143, 109906 (2020).

160. Gould, T. J. et al. Neutrophil extracellular traps promote thrombin generation through platelet-dependent and platelet-independent mechanisms. Arterioscler. Thromb. Vasc. Biol. 34, 1977–1984 (2014).

161. Wang, Y. et al. Neutrophil extracellular trap-microparticle complexes enhance thrombin generation via the intrinsic pathway of coagulation in mice. Sci. Rep. 8, 1–14 (2018).

162. Zuo, Y. et al. Neutrophil extracellular traps and thrombosis in COVID-19. J. Thromb. Thrombolysis 51, 446–453 (2021).

163. Hisada, Y. et al. Neutrophils and neutrophil extracellular traps enhance venous thrombosis in mice bearing human pancreatic tumors. Haematologica 105, 218–225 (2020).

164. Karki, R. et al. Synergism of TNF-α and IFN-γ Triggers Inflammatory Cell Death, Tissue Damage, and Mortality in SARS-CoV-2 Infection and Cytokine Shock Syndromes. Cell 184, 149–168.e17 (2021).

165. Eloseily, E. M. et al. Benefit of Anakinra in Treating Pediatric Secondary Hemophagocytic Lymphohistiocytosis. Arthritis Rheumatol. 72, 326–334 (2020).

166. Durand, M., Troyanov, Y., Laflamme, P. & Gregoire, G. Macrophage activation syndrome treated with anakinra. J. Rheumatol. 37, 879–880 (2010).

167. Kang, S., Tanaka, T., Narazaki, M. & Kishimoto, T. Targeting Interleukin-6 Signaling in Clinic. Immunity 50, 1007–1023 (2019).

168. van der Stegen, S. J. C. et al. Preclinical In Vivo Modeling of Cytokine Release Syndrome Induced by ErbB-Retargeted Human T Cells: Identifying a Window of Therapeutic Opportunity? J. Immunol. 191, 4589–4598 (2013).

169. Teachey, D. T. et al. Cytokine release syndrome after blinatumomab treatment related to abnormal macrophage activation and ameliorated with cytokine-directed therapy. Blood 121, 5154–5157 (2013).

170. Winkler, U. et al. Cytokine-release syndrome in patients with B-cell chronic lymphocytic leukemia and high lymphocyte counts after treatment with an anti-CD20 monoclonal antibody (rituximab, IDEC-C2B8). Blood 94, 2217–2224 (1999).

171. Faulkner, L., Cooper, A., Fantino, C., Altmann, D. M. & Sriskandan, S. The Mechanism of Superantigen-Mediated Toxic Shock: Not a Simple Th1 Cytokine Storm. J. Immunol. 175, 6870–6877 (2005).

172. Ablamunits, V. & Lepsy, C. Blocking TNF signaling may save lives in COVID-19 infection. Mol. Biol. Rep. 49, 2303–2309 (2022).

173. Guo, Y. et al. Targeting TNF-α for COVID-19: Recent Advanced and Controversies. Front. Public Heal. 10, 1–9 (2022).

174. Saraiva, M. et al. Biology and therapeutic potential of interleukin-10. J. Exp. Med. 217, 1–19 (2020).

175. Fioranelli, M. & Roccia, M. G. Twenty-five years of studies and trials for the therapeutic application of IL-10 immunomodulating properties. From high doses administration to low dose medicine new paradigm. J. Integr. Cardiol. 1, 2–6 (2014).

176. Kircheis, R. et al. NF-κB Pathway as a Potential Target for Treatment of Critical Stage COVID-19 Patients. Front. Immunol. 11, 1–11 (2020).

177. Coldewey, S. M., Rogazzo, M., Collino, M., Patel, N. S. A. & Thiemermann, C. Inhibition of IκB kinase reduces the multiple organ dysfunction caused by sepsis in the mouse. DMM Dis. Model. Mech. 6, 1031–1042 (2013).

178. Yamamoto, Y. & Gaynor, R. B. Therapeutic potential of inhibition of the NF-κB pathway in the treatment of inflammation and cancer. J. Clin. Invest. 107, 135–142 (2001).

179. Ponta, H., Kanno, T., Franzoso, G., Helmberg, A. & Karin, M. GR could physically associate with NF-KB Immunosuppression by Glucocorticoids: Inhibition of NF-KB Activity Through IKB Synthesis. Science (80-.). 270, 286–290 (1993).

180. Fan, P. et al. Suppression of nuclear factor-kB by glucocorticoid receptor blocks estrogen-induced apoptosis in estrogen-deprived breast cancer cells. Mol. Cancer Ther. 18, 1684–1695 (2019).

181. McComb, S. et al. Type-I interferon signaling through ISGF3 complex is required for sustained Rip3 activation and necroptosis in macrophages. Proc. Natl. Acad. Sci. U. S. A. 111, 3206–3213 (2014).

182. Yanai, H. et al. Revisiting the role of IRF3 in inflammation and immunity by conditional and specifically targeted gene ablation in mice. Proc. Natl. Acad. Sci. U. S. A. 115, 5253–5258 (2018).

183. Xu, X., Wang, W., Lin, L. & Chen, P. Liraglutide in combination with human umbilical cord mesenchymal stem cell could improve liver lesions by modulating TLR4/NF-kB inflammatory pathway and oxidative stress in T2DM/NAFLD rats. Tissue Cell 66, 101382 (2020).

184. Jiang, Z. & Zhang, J. Mesenchymal stem cell-derived exosomes containing miR-145-5p reduce inflammation in spinal cord injury by regulating the TLR4/NF-κB signaling pathway. Cell Cycle 20, 993–1009 (2021).

185. Su, V. Y. F., Lin, C. S., Hung, S. C. & Yang, K. Y. Mesenchymal stem cell-conditioned medium induces neutrophil apoptosis associated with inhibition of the NF-κb pathway in endotoxin-induced acute lung injury. Int. J. Mol. Sci. 20, (2019).

186. Liu, Y. et al. Human placental mesenchymal stem cells regulate inflammation via the NF-κB signaling pathway. Exp. Ther. Med. 24, 1–11 (2022).

187. Yang, X. et al. Flagellin attenuates experimental sepsis in a macrophage-dependent manner. Crit. Care 23, 1–14 (2019).

188. Belikoff, B. G. et al. A2B Adenosine Receptor Blockade Enhances Macrophage-Mediated Bacterial Phagocytosis and Improves Polymicrobial Sepsis Survival in Mice. J. Immunol. 186, 2444–2453 (2011).

189. Cui, J. et al. The clinical efficacy of intravenous IgM-enriched immunoglobulin (pentaglobin) in sepsis or septic shock: a meta-analysis with trial sequential analysis. Ann. Intensive Care 9, (2019).

190. Busani, S., Damiani, E., Cavazzuti, I., Donati, A. & Girardis, M. Intravenous immunoglobulin in septic shock: Review of the mechanisms of action and meta-analysis of the clinical effectiveness. Minerva Anestesiol. 82, 559–572 (2016).

191. Akdag, A. et al. Role of Pentoxifylline and/or IgM-Enriched Intravenous Immunoglobulin in the Management of Neonatal Sepsis. Am. J. Perinatol. 31, 905–912 (2014).

192. Greenfield, K. G., Badovinac, V. P., Griffith, T. S. & Knoop, K. A. Sepsis, Cytokine Storms, and Immunopathology: The Divide between Neonates and Adults. ImmunoHorizons 5, 512–522 (2021).

193. Kuhn, P. et al. A Multicenter, Randomized, Placebo-Controlled Trial of Prophylactic Recombinant Granulocyte-Colony Stimulating Factor in Preterm Neonates with Neutropenia. J. Pediatr. 155, (2009).

194. Miura, E., Procianoy, R. S., Bittar, C. & Miura, C. S. With the Clinical Diagnosis of Early-Onset Sepsis. 107, (2015).

195. Bo, L., Wang, F., Zhu, J., Li, J. & Deng, X. Granulocyte-colony stimulating factor (G-CSF) and granulocyte-macrophage colony stimulating factor (GM-CSF) for sepsis: A meta-analysis. Crit. Care 15, 1–12 (2011).

196. Mathias, B., Szpila, B. E., Moore, F. A., Efron, P. A. & Moldawer, L. L. A review of GM-CSF therapy in sepsis. Med. (United States) 94, 1–10 (2015).

197. Pool, R., Gomez, H. & Kellum, J. A. Mechanisms of Organ Dysfunction in Sepsis. Crit. Care Clin. 34, 63–80 (2018).

198. Spapen, H. D., Jacobs, R. & Honoré, P. M. Sepsis-induced multi-organ dysfunction syndrome—a mechanistic approach. J. Emerg. Crit. Care Med. 1, 27–27 (2017).

199. Tamburro, R. F. & Jenkins, T. L. Multiple organ dysfunction syndrome: A challenge for the pediatric critical care community. Pediatr. Crit. Care Med. 18, S1–S3 (2017).

200. Wang, H. & Ma, S. The cytokine storm and factors determining the sequence and severity of organ dysfunction in multiple organ dysfunction syndrome. Am. J. Emerg. Med. 26, 711–715 (2008).

201. Lelubre, C. & Vincent, J. L. Mechanisms and treatment of organ failure in sepsis. Nat. Rev. Nephrol. 14, 417–427 (2018).

