## supplemental figs for "Mesenchymal stromal cells and alpha-1 antitrypsin have a strong synergy in modulating inflammation and its resolution"

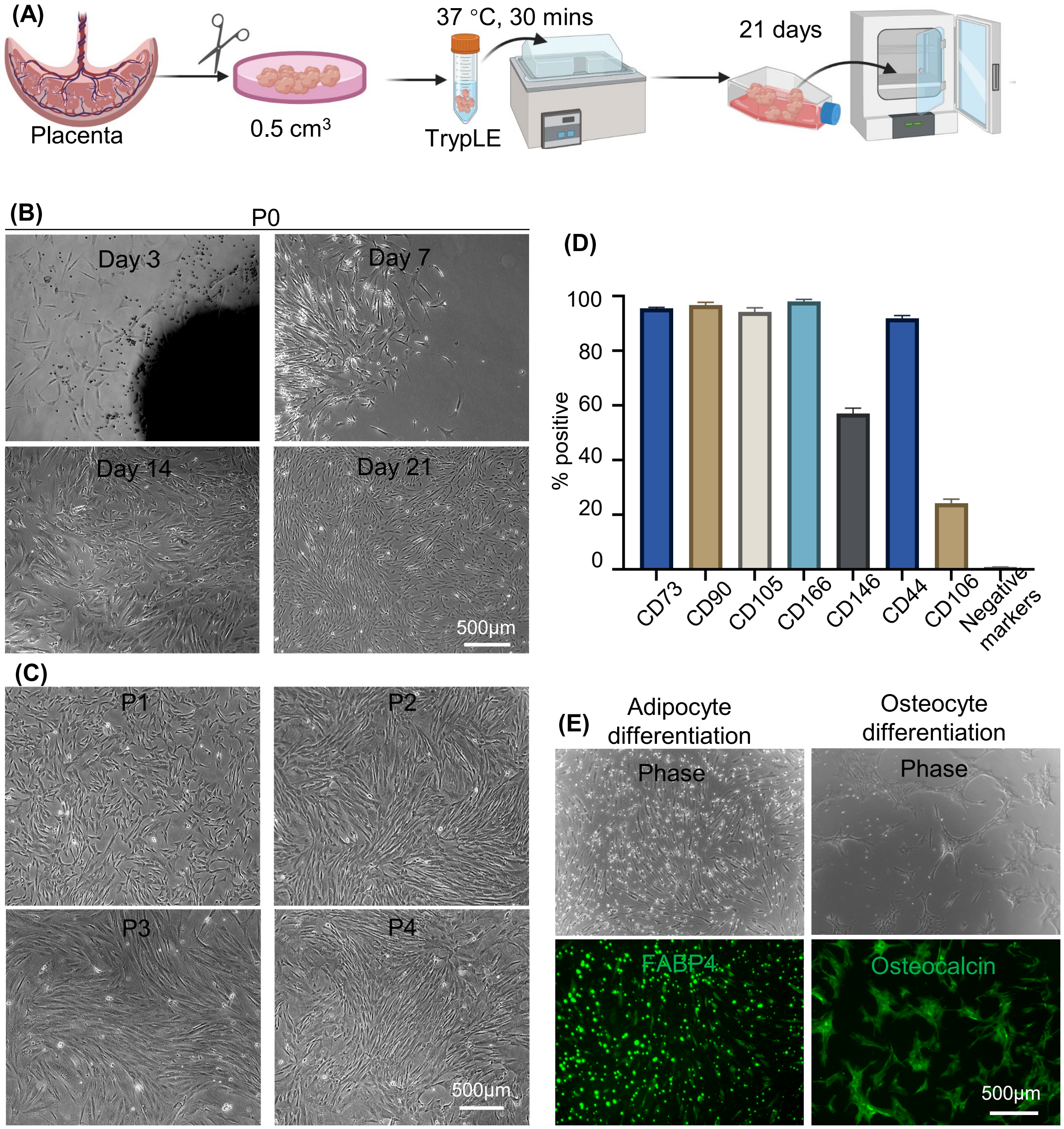


**Fig.S1** (A) Mesenchymal stromal cells (MSCs) isolation process. (B) MSCs migrating from a small tissue. (C) MSCs during the expansion phase (Passage 1 to 4). (D) Surface marker expression for P4 MSCs. Negative markers include CD34, CD45, CD11b, CD79A, and HLA-DR. (E) P4 MSCs could be differentiated into FABP4+ adipocytes and osteocalcin+ osteocytes.

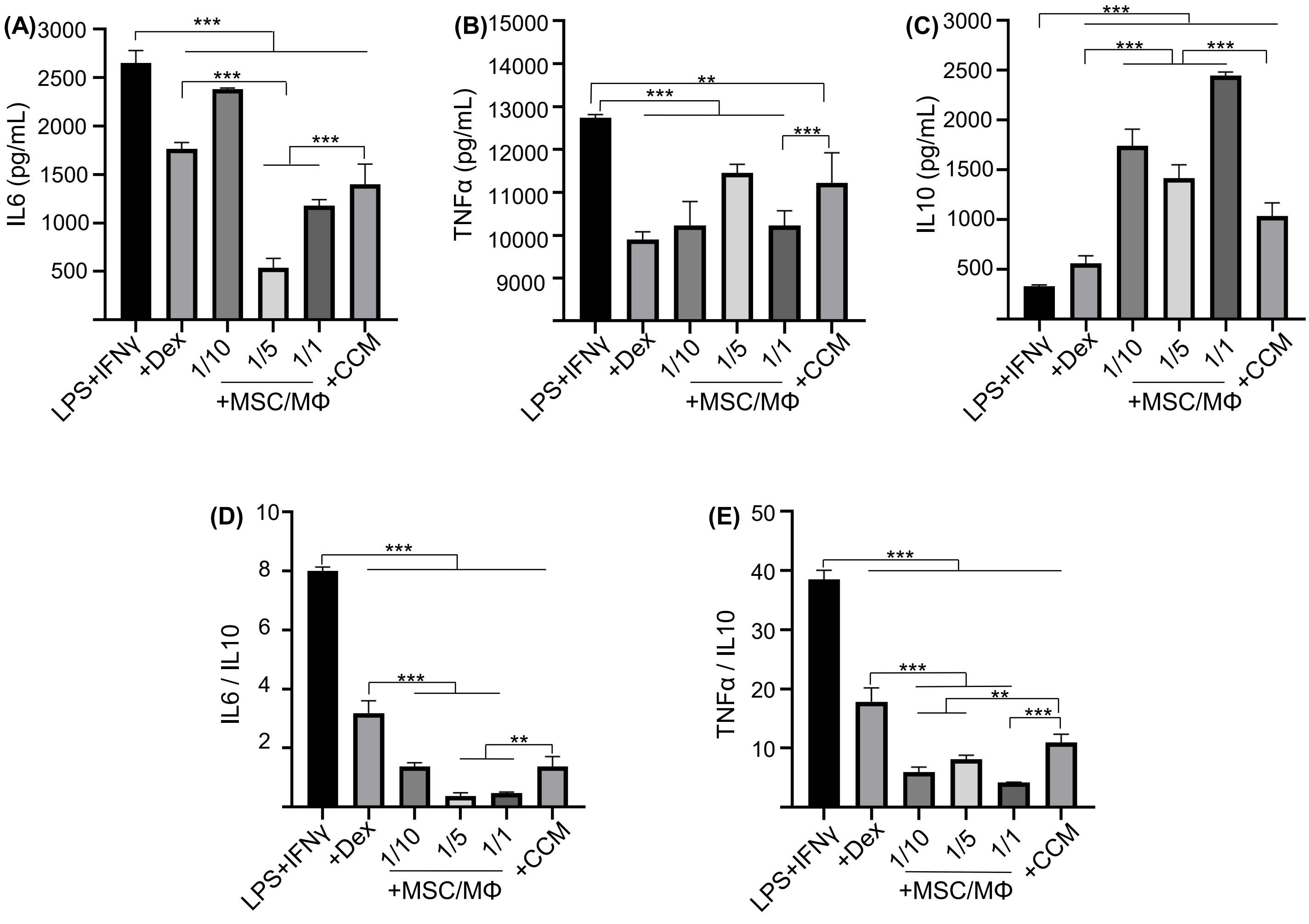


**Fig.S2** MSCs modulated inflammation in Raw 264.7 macrophages (MΦs). Cells were stimulated with 100 ng/mL LPS plus 10 ng/mL IFNγ and co-cultured with MSCs at different ratios (MSC/MΦ=1/10, or 1/5 or 1/1) or treated with MSC conditioned medium (CCM). Dexamethasone (Dex, 1 μg/mL) was used as a benchmark. Pro-inflammatory mouse cytokine IL6 (A) TNFα (B) and anti-inflammatory mouse cytokine IL10 (C) were measured via ELISA. The IL6/IL10 (D) and TNFα/IL10 ratio (E) were also shown. **:p<0.05, **:p<0.01, ***:p<0.001.*

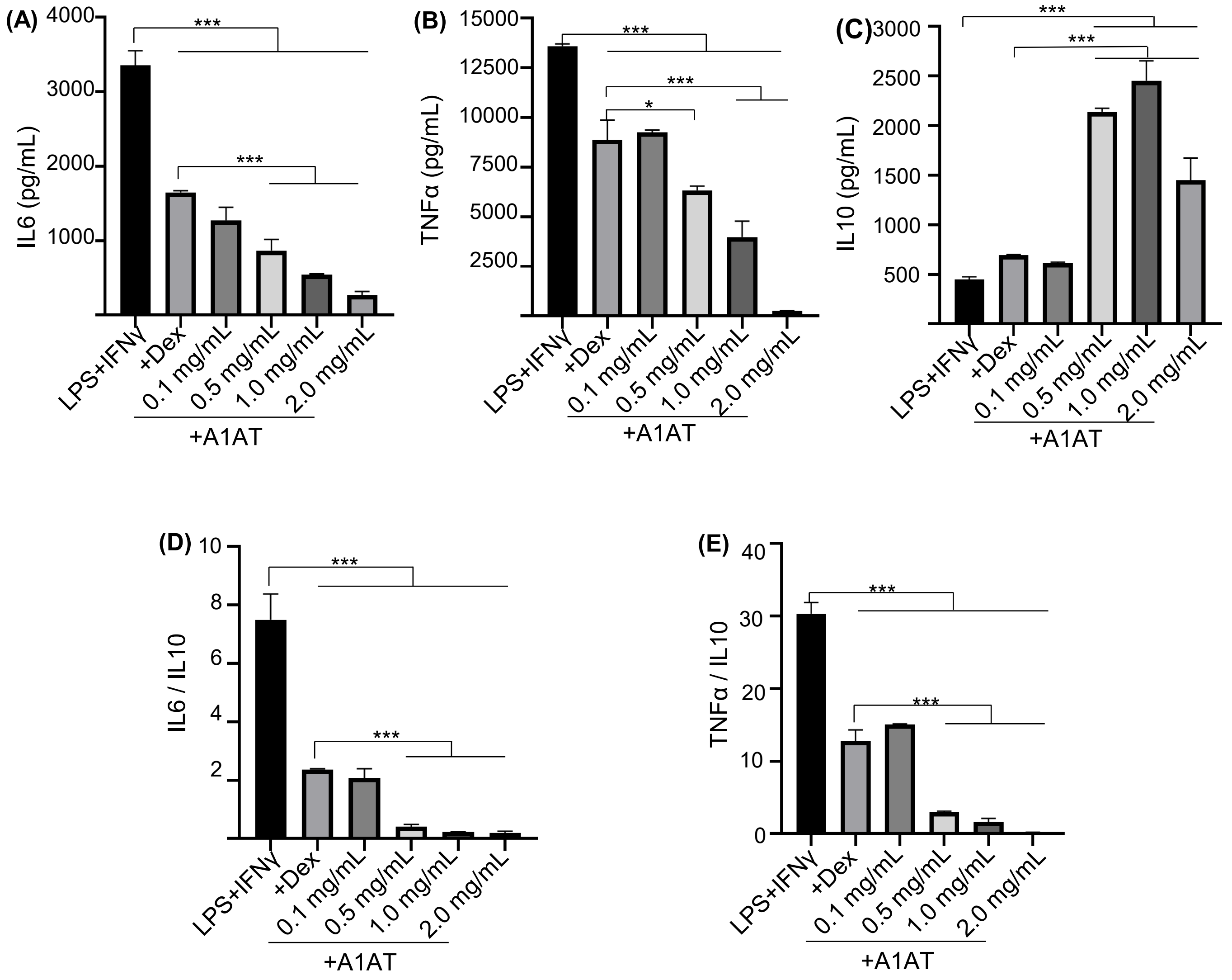


**Fig.S3** A1AT modulated inflammation in Raw 264.7 macrophages. Cells were stimulated with 100 ng/mL LPS plus 10 ng/mL IFNγ and treated with A1AT at 0.1 to 2.0 mg/mL. Dexamethasone (Dex, 1 μg/mL) was used as benchmark. Pro-inflammatory mouse cytokine IL6 (A), TNFα (B) and anti-inflammatory mouse cytokine IL10 (C) were measured via ELISA. The IL6/IL10 (D) and TNFα/IL10 ratio (E) were also shown. **:p<0.05, **:p<0.01, ***:p<0.001.*

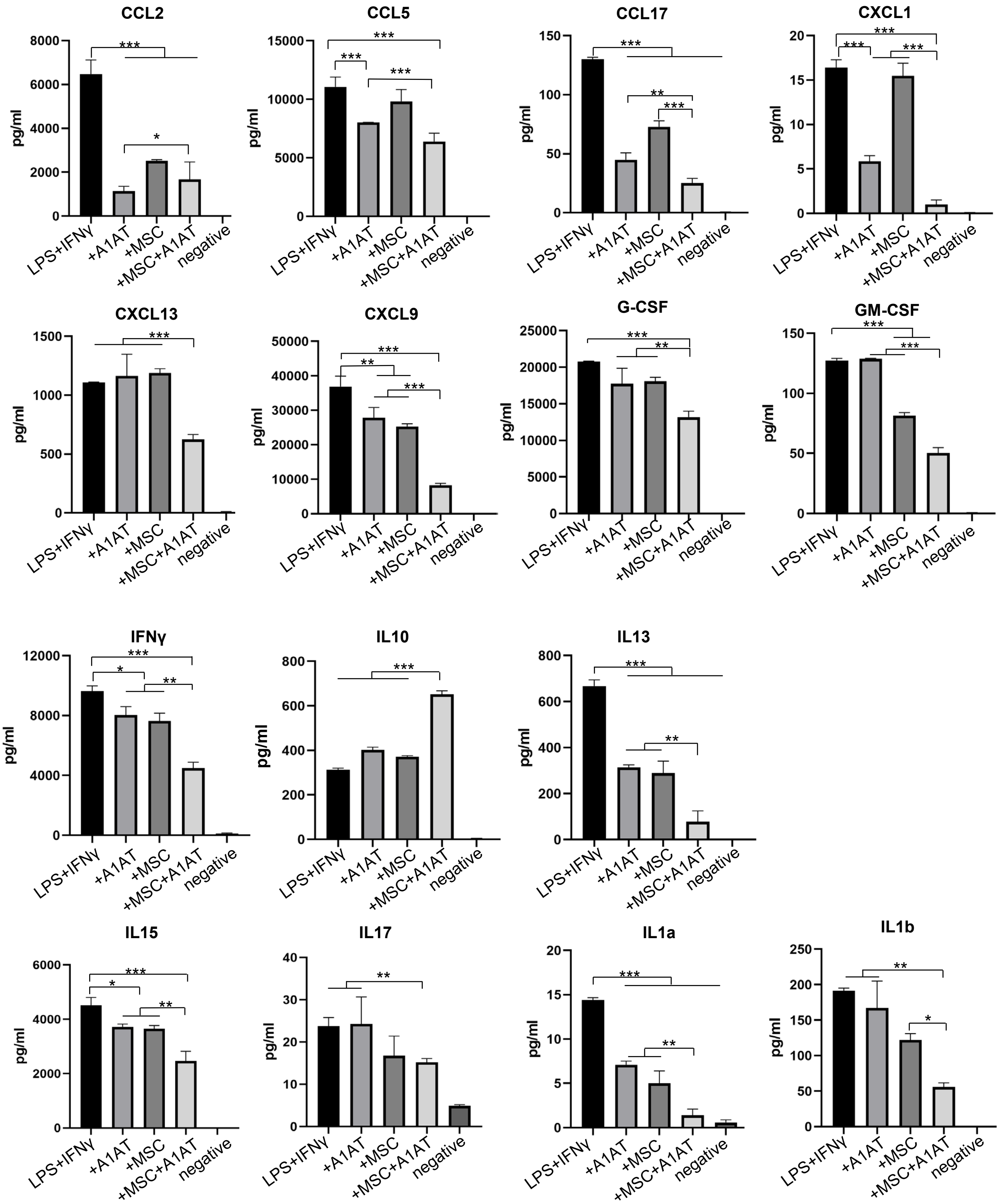
CCL2
CCL5
CCL17
CXCL1
CXCL13
CXCL9
G-CSF
GM-CSF
IFNγ
IL10
IL13
IL15
IL17
IL1a
IL1b
pg/ml
LPS+IFNγ
+A1AT
+MSC
+MSC+A1AT
negative

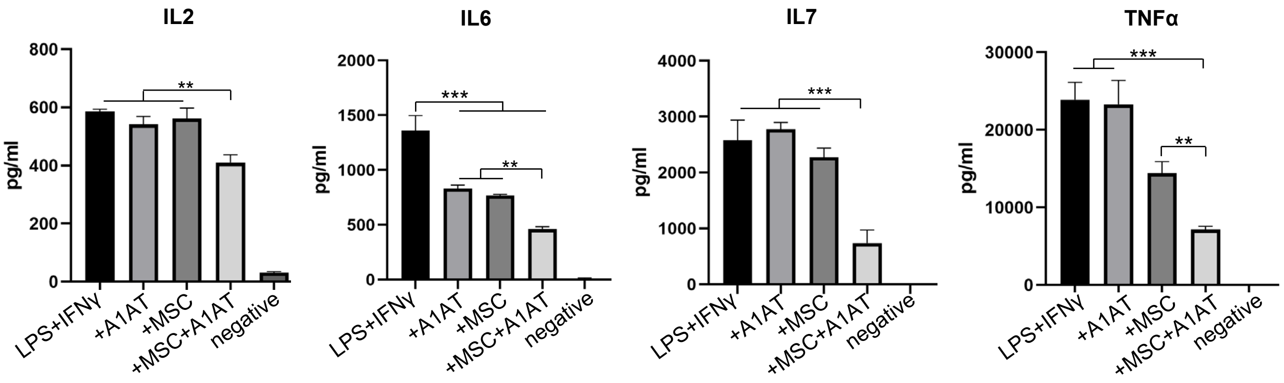


**Fig.S4** MSCs synergized with A1AT to modulate inflammation in Raw 264.7 macrophages. Cells were stimulated with 100 ng/mL LPS plus 10 ng/mL IFNγ and treated with 0.5 mg/mL A1AT or MSCs (MSC/MΦ=1/10) or their combination. Dexamethasone (Dex, 1 μg/mL) was used as a benchmark. 40 mouse cytokines in the medium were measured. Negative: cells were not stimulated and had no treatment. **:p<0.05, **:p<0.01, ***:p<0.001.*

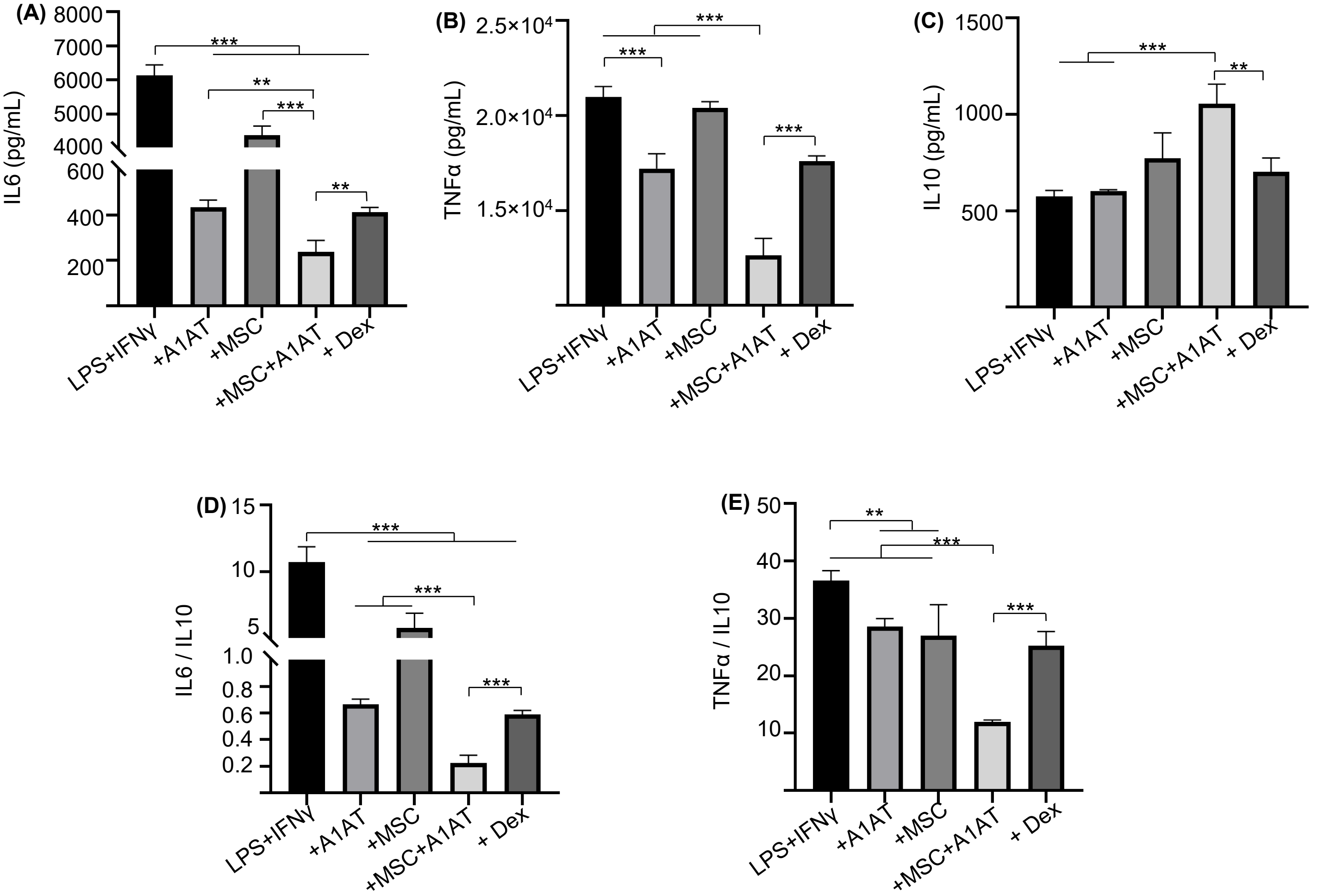


**Fig.S5** MSCs synergized with A1AT to modulate inflammation in human THP-1 monocytes derived macrophages. Cells were stimulated with 100 ng/mL LPS plus 10 ng/mL IFNγ and treated with 0.5 mg/mL A1AT or MSCs (MSC/MΦ=1/10) or their combination. Dexamethasone (Dex, 1 µg/mL) was used as a benchmark. Pro-inflammatory human cytokine IL6 (A), TNFα (B) and anti-inflammatory human cytokine IL10 (C) were measured via ELISA. The IL6/IL10 (D) and TNFα/IL10 ratio (E) were also shown. **:p<0.05, **:p<0.01, ***:p<0.001.*

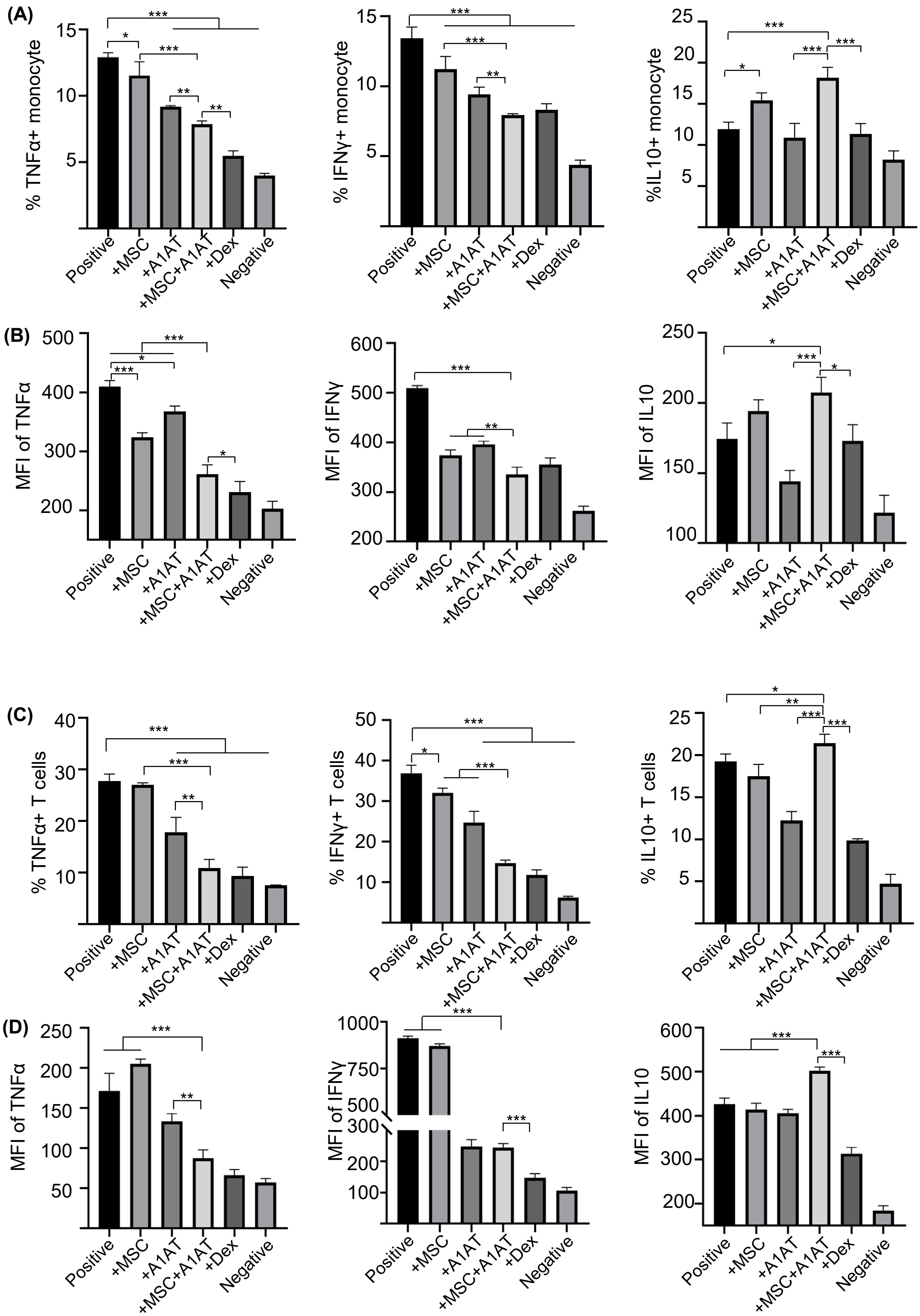


**Fig. S6** MSCs synergized with A1AT to modulate inflammation in primary human PBMCs. Cells were stimulated with 100 ng/mL LPS + anti-CD3/CD28 antibodies (positive) and treated with 0.5 mg/mL A1AT or MSCs (MSC/PBMC=1/10) or their combination for 72 hrs. Dexamethasone (Dex, 1 µg/mL) was used as a benchmark. PBMCs without activation and treatment was used as a negative control. (A) The % TNFα, IFNγ and IL10 positive monocytes. (B) The mean fluorescence intensity (MFI) monocyte. (C) The % TNFα, IFNγ and IL10 positive T cells and (D) the MFI of TNFα, IFNγ and IL10 per T cell. **:p<0.05, **:p<0.01, ***:p<0.001.*

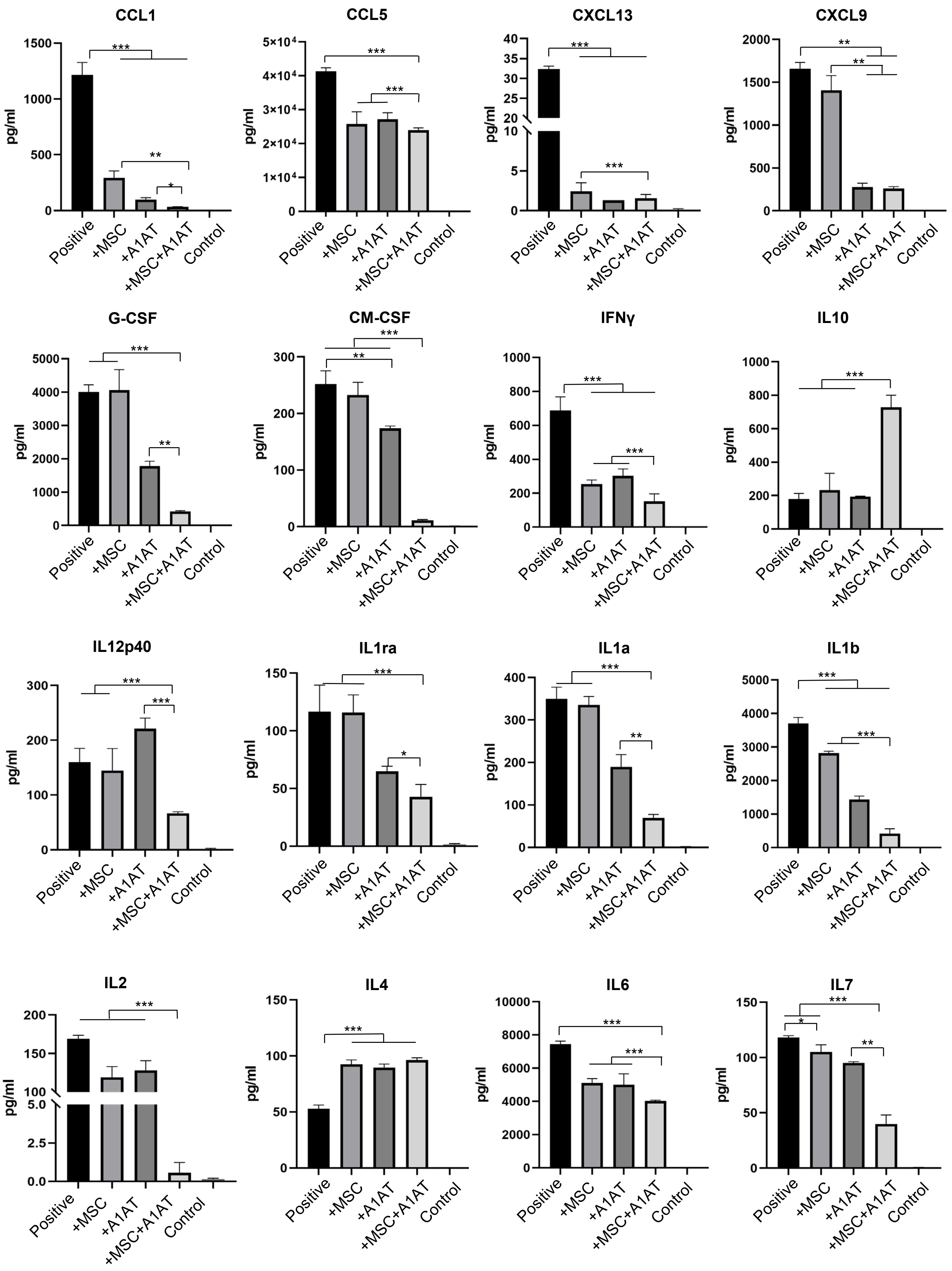
CCL1
CCL5
CXCL13
CXCL9
G-CSF
CM-CSF
IFNγ
IL10
IL12p40
IL1ra
IL1a
IL1b
IL2
IL4
IL6
IL7
pg/ml
Positive
+MSC
+A1AT
+MSC+A1AT
Control
*
**
***

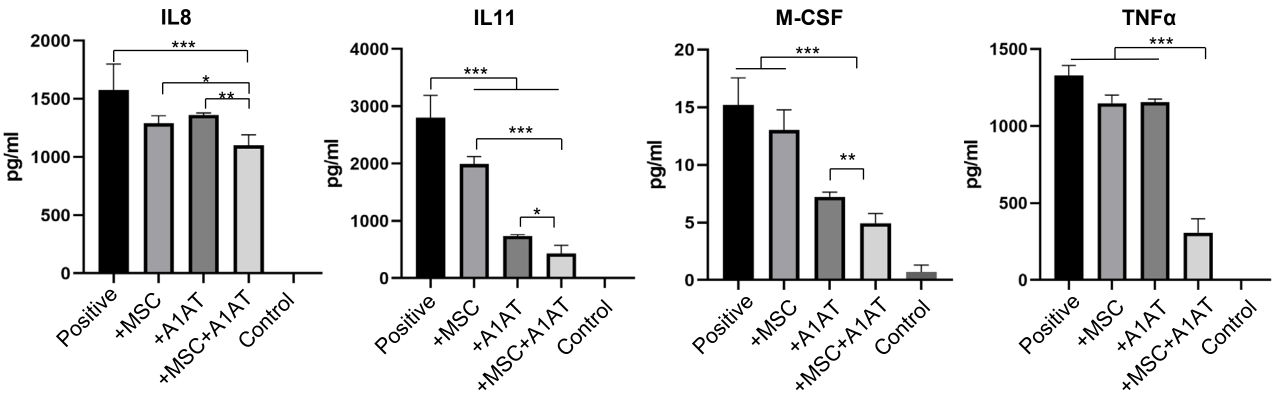


**Fig.S7** MSCs synergized with A1AT to modulate inflammation in primary human PBMCs. Cells were stimulated with 100 ng/mL LPS + anti-CD3/CD28 antibodies (positive) and treated with 0.5 mg/mL A1AT or MSCs (MSC/PBMC=1/10) or their combination for 24 hrs. Dexamethasone (Dex, 1 μg/mL) was used as a benchmark. 40 human cytokines in the medium were measured. Negative: cells were not stimulated and had no treatment. **:p<0.05, **:p<0.01, ***:p<0.001.*

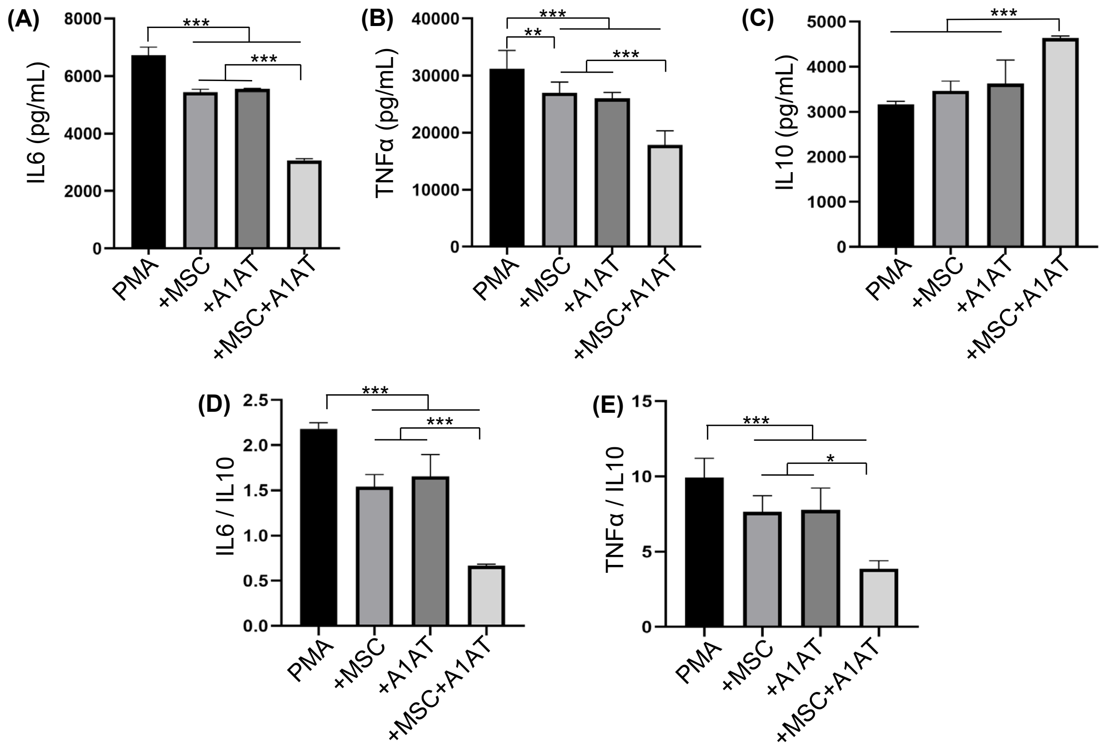


**Fig.S8** MSCs synergized with A1AT to modulate inflammation in neutrophils. HL-60 cells were differentiated into neutrophils with DMSO (1.25% v/v) and ATRA (0.1 μM) for 3 days. Neutrophils were stimulated with 100 nM PMA and and treated with 0.5 mg/mL A1AT or MSCs (MSC/neutrophil=1/10) or their combination. Pro-inflammatory human cytokine IL6 (A), TNFα (B) and anti-inflammatory human cytokine IL10 (C) were measured via ELISA. The IL6/IL10 (D) and TNFα/IL10 ratio (E) were also shown. **:p<0.05, **:p<0.01, ***:p<0.001.*

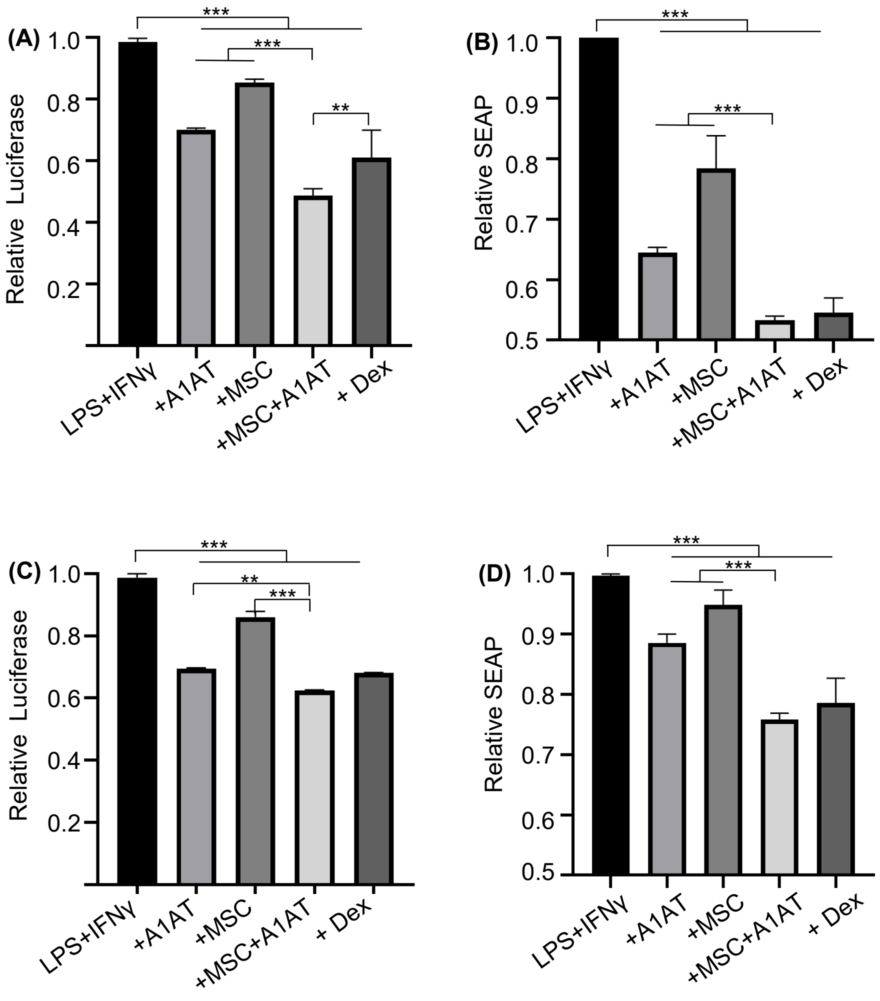


**Fig.S9** MSCs synergized with A1AT to modulate NF-κB and IRFs signaling. Raw 264.7 (A, B) and THP-1 derived macrophages (C, D) expressing a luciferase reporter for IRFs signaling and a secreted alkaline phosphatase (SEAP) reporter for NF-κB signaling. Cells were stimulated with 100 ng/mL LPS plus 10 ng/mL IFN-γ and treated with 0.5 mg/mL A1AT or MSCs (MSC/MΦ=1/10) or their combination. Dexamethasone (Dex, 1 μg/mL) was used as a benchmark. Luciferase (A, C) and SEAP (B, D) activities were quantified. **:p<0.05, **:p<0.01, ***:p<0.001.*
